# Phosphorylation-dependent routing of RLP44 towards brassinosteroid or phytosulfokine signalling

**DOI:** 10.1101/527754

**Authors:** Borja Garnelo Gómez, Eleonore Holzwart, Andreas Kolbeck, Chaonan Shi, Rosa Lozano-Durán, Sebastian Wolf

## Abstract

Plants rely on a complex network of cell surface receptors to integrate developmental and environmental cues into behaviour adapted to the conditions. The largest group of these receptors, leucine-rich repeat receptor-like kinases, form a complex interaction network that is modulated and extended by receptor-like proteins. This raises the question of how specific outputs can be generated when receptor proteins are engaged in a plethora of promiscuous interactions. RECEPTOR-LIKE PROTEIN 44 (RLP44) acts to promote both brassinosteroid and phytosulfokine signalling, which orchestrate a wide variety of cellular responses. However, it is unclear how these activities are coordinated. Here, we show that RLP44 is phosphorylated in its highly conserved C-terminal cytosolic tail and that this post-translational modification governs its subcellular localization. RLP44 variants in which phosphorylation is blocked enter endocytosis prematurely, leading to an almost entirely intracellular localization, whereas phospho-mimicking or ectopic phosphorylation results in preferential RLP44 localization at the plasma membrane. Phosphorylation of the C-terminus is essential for brassinosteroid-associated functions of RLP44. In contrast, RLP44’s role in phytosulfokine signalling is not affected by its phospho-status. Detailed mutational analysis suggests that phospho-charge, rather than modification of individual amino acids determines routing of RLP44 to its target receptor complexes, providing a framework to understand how a common component of different receptor complexes can get specifically engaged in a particular signalling pathway.

## Introduction

To integrate environmental cues with intrinsic developmental programs, plants depend on the perception of extracellular signals by their expanded family of cell surface receptors. The largest group of these plasma membrane-localized proteins is formed by leucine-rich repeat receptor-like kinases (LRR-RLKs), characterized by the presence of a signal peptide for entry into the secretory pathway, a leucine-rich repeat-containing extracellular domain, a single pass transmembrane domain, and an intracellular kinase domain with homology to animal Irak/Pelle proteins (Shiu and Bleecker, 2001). LRR-RLKs form an extensive interaction network reflecting the need to decipher complex information about the environment (Smakowska-Luzan et al., 2018). LRR-RLPs, which resemble LRR-RLKs, but lack a kinase domain, contribute to RLK signalling in a variety of ways (Jamieson et al., 2018), further expanding the complexity of the network. Thus, the multitude of potential interactions and extensive sharing of components between pathways, raise a central question in plant signal transduction: how can distinct signalling responses be achieved? A partial answer is provided by the discovery of membrane sub-compartmentalization, which helps to spatially separate potential interaction partners (Gronnier et al., 2019; Jaillais and Ott, 2019; Galindo-Trigo et al., 2020). Furthermore, selective endocytosis, recycling, and eventually, the degradation of RLKs in the vacuole (Martins et al., 2015; Mbengue et al., 2016; Zhou et al., 2018; Liu et al., 2020) are crucial for tuning of signalling. Most, if not all plasma membrane receptors described so far undergo endocytosis in vesicles coated by clathrin, which depend on cytosolic adaptor complexes for cargo selection and transport (Van Damme et al., 2011; Di Rubbo et al., 2013; Gadeyne et al., 2014; Liu et al., 2020). In some, but not all, cases, ligand binding promotes internalization, possibly to establish a refractory phase after ligand exposure and prevent potentially harmful continuous activation (Robatzek et al., 2006; Nimchuk et al., 2011; Ortiz-Morea et al., 2016). Endocytic trafficking of LRR-RLK proteins is intimately linked to post-translational modification of their cytosolic domains through addition of the small protein ubiquitin to lysine residues (Dubeaux and Vert, 2017). A different kind of post-translational modification, phosphorylation, is at the core of RLK signalling regulation. Binding of extracellular ligands to the ectodomains of several LRR-RLKs has been shown to mediate hetero-dimerization with a shape-complementary co-receptor of the SOMATIC-EMBRYOGENESIS RECEPTOR KINASE (SERK) family (Albert et al., 2013; Couto and Zipfel, 2016; Hohmann et al., 2017; Mithoe and Menke, 2018), juxtaposing their kinase domains in the cytosol. This, in turn, leads to auto and trans-phosphorylation of the kinases, resulting in an activated receptor complex capable of recruiting and phosphorylating downstream signalling components. Recent evidence suggests differential requirements for individual phosphorylation events in SERK kinase domains depending on the interacting RLK (Perraki et al., 2018). Signalling output is further modified by the activity of phosphatases and the phosphorylation-dependent release of inhibitory factors (Park et al., 2008; Jaillais et al., 2011b; Lin et al., 2013; Monaghan et al., 2014; Couto et al., 2016). In summary, post-translational modification through phosphorylation and ubiquitination are key mechanisms to spatially and temporally control LRR-RLK-mediated signalling. One of the best-characterized LRR-RLKs is the receptor for brassinosteroid (BR) phytohormones, BRASSINOSTEROID INSENSITIVE 1 (BRI1) (Li and Chory, 1997). Signalling mediated by BRI1 and its co-receptors, such as SOMATIC-EMBRYOGENESIS RECEPTOR KINASE 1/BRI1-ASSOCIATED KINASE1 (SERK3/BAK1) (Li et al., 2002; Nam and Li, 2002), plays a crucial role in cell elongation, in part by controlling a plethora of cell wall biosynthesis and remodelling genes (Sun et al., 2010; Belkhadir and Jaillais, 2015; Singh and Savaldi-Goldstein, 2015). We previously revealed that when cell wall integrity is challenged by interference with the activity of the important cell wall modification enzyme pectin methylesterase (PME), BR signalling is activated as a compensatory mechanism (Wolf et al., 2012b). This BR-mediated compensatory response depends on RECEPTOR-LIKE PROTEIN 44 (RLP44) (Wolf et al., 2014). RLP44 directly interacts with both BRI1 and BAK1 and promotes their association (Holzwart et al., 2018). However, RLP44 also interacts with and promotes the activity of the receptor complex for the plant growth peptide phytosulfokine (PSK) (Holzwart et al., 2018, 2020; Sauter, 2015). The interaction between RLP44 and the PSK receptor PSKR1 is important for the maintenance of procambial cell fate, as both RLP44 and PSK-related mutants show ectopic xylem formation in the position of the procambium in seedling roots (Holzwart et al., 2018, 2020). Thus, RLP44 acts in at least two different LRR-RLK pathways through direct interaction with their receptors. Multi-faceted interactions among LRR proteins is an emerging theme in plant receptor biology (Ma et al., 2016; Smakowska-Luzan et al., 2018); however, it is not clear how distinct responses are ensured. As RLP44 acts in two separate pathways with well-defined read-outs, it provides an excellent model to decipher how pathway specificity is achieved. Here, we show that RLP44 is phosphorylated in its highly conserved C-terminal cytosolic tail. This post-translational modification is crucial for regulating RLP44’s function in BR signalling activation. RLP44 variants in which phosphorylation is blocked enter endocytosis prematurely, leading to an almost entirely intracellular localization. Conversely, mimicking phosphorylation or ectopic phosphorylation results in preferential RLP44 localization at the plasma membrane. This increase in the ratio of plasma membrane to intracellular localization is controlled by phospho-charge, rather than by modification of specific amino acids and is furthermore dependent on the presence of BRI1, suggesting that phosphorylation affects subcellular localization through modulating the interactions of the LRR proteins. In contrast, RLP44’s role in PSK signalling is not affected by phospho-status. Thus, our results provide a framework to understand how specificity can be determined in plasma membrane receptor complex interactions.

## Results

### Four conserved putative phosphorylation sites are required for RLP44-mediated BR signalling activation

RLP44 is unusual compared to other RLPs in Arabidopsis, as its juxtamembrane domain is not acidic, and its cytoplasmic, C-terminal tail shows a pI of 4.7, whereas the majority of Arabidopsis RLPs harbour cytoplasmic tails with basic pI (Gust and Felix, 2014). However, this unusual cytoplasmic domain (CD) is well conserved among the apparent RLP44 orthologues in land plants (Figure 1A and Supplemental Figure S1). Interestingly, of four putative phosphorylation sites in AtRLP44 (from hereon RLP44), three - T256, S268, and Y274 - are conserved in all orthologues, whereas S270 seems to be specific to *Brassicacea*. All four sites are predicted to be phosphorylated (NetPhos 3.1 server (www.cbs.dtu.dk/services/NetPhos/)) and we have previously obtained evidence for serine phosphorylation in RLP44 using anti-phosphoserine antibodies (Wolf et al., 2014). To assess which of the four amino acids are phosphorylated in vivo, we performed mass spectroscopy after immunoprecipitation from transgenic Arabidopsis plants expressing an RLP44-GFP fusion protein under control of the CaMV35S promoter, as well as from transiently transformed *Nicotiana benthamiana* leaves. Peptide coverage was quite poor, in particular from Arabidopsis, despite effective immune-purification of the RLP44 fusion protein. However, we were able to identify S268 phosphorylation in *N. benthamiana* (Supplemental Figure S2). As we could not rule out modification of the other three residues, we first assessed the effect of blocking post-translational modification of all four putative phospho-sites. To this end, we generated a version of RLP44 fused to GFP in which all four sites are mutated to either alanine (T256A, S268A, A270A) or phenylalanine (Y274F) and termed this “phospho-dead” variant RLP44-GFP Pdead. Conversely, we created a “phospho-mimic” (Pmimic) version of RLP44-GFP in which all four putative phospho-sites are converted to Glutamate (T256E, S286E, A270E, Y274E). Throughout the manuscript, we refer to these genetic modifications of putative phosphorylation sites as affecting “phospho-status” for brevity. Constructs encoding these mutant versions of RLP44-GFP along with a wild-type version (RLP44-GFP WT) were used to transform the *rlp44* mutant *cnu2* (Wolf et al., 2014) for complementation assays. We originally described *cnu2* as a suppressor mutant of an overexpression line of *PMEI5*(PMEIox), which displays growth defects due to a compensatory boosting of brassinosteroid signalling strength. Activation of BR signalling in response to PMEI-mediated reduction of PME activity critically depends on RLP44, which directly interacts with the BR receptor BRI1 and its co-receptor BAK1. Mutation of RLP44 in *cnu2* thus leads to relatively normal growth despite the presence of the PMEIox transgene. As previously described (Wolf et al., 2014), expression of RLP44-GFP is able to complement *cnu2* and results in recovery of the PMEIox seedling root waving phenotype in several independent transgenic lines (Figure 1C and Supplemental Figure S3A). Similarly, the Pmimic variant of RLP44-GFP could restore or even slightly enhance the PMEIox phenotype, suggesting that the presence of the native version of the four mutated RLP44 sites is not essential for function in BR signalling activation, and that mimicking phosphorylation at these sites might be associated with enhanced activity. In contrast, the Pdead version consistently failed to complement *cnu2* with respect to the root waving phenotype (Figure 1D and Supplemental Figure S3A). Concentrating on one line for each construct with comparable RLP44-GFP expression levels (Supplemental Figure S3B), we made similar observations for other previously described PMEIox phenotypes (Wolf et al., 2012b), such as altered expression of BR marker genes (Supplemental Figure S3C, D), reduced seed yield (Supplemental Figure S3E), and agravitropic growth on vertical agar plates in the dark due to enhanced BR signalling (Supplemental Figure S4). In each case, the line expressing the WT or Pmimic version of RLP44-GFP in the *cnu2* suppressor mutant behaved like PMEIox, whereas the line expressing the Pdead version behaved like *cnu2*, i.e. similar to wild type. We then assessed the ability of the three RLP44-GFP versions to rescue *rlp44* phenotypes in the absence of PMEIox-induced cell wall challenge. Mutants of RLP44 such as *rlp44^cnu2^* show reduced petiole length (Wolf et al., 2014), presumably caused by impaired BR signalling. Expression of RLP44-GFP WT and RLP4-GFP Pmimic, but not RLP44-GFP Pdead, could restore the petiole length defect of *rlp44^cnu2^* (Figure 2), in line with the assumption that this phenotype is BR signalling-related. In conclusion, blocking phosphorylation of four putative phosphosites in the cytoplasmic domain precluded BR signalling activation-related functions of RLP44. In contrast, introducing a negative charge to mimic phosphorylation at these sites resulted in functionality comparable to that of the wild-type version of RLP44-GFP.

**Figure 1.**
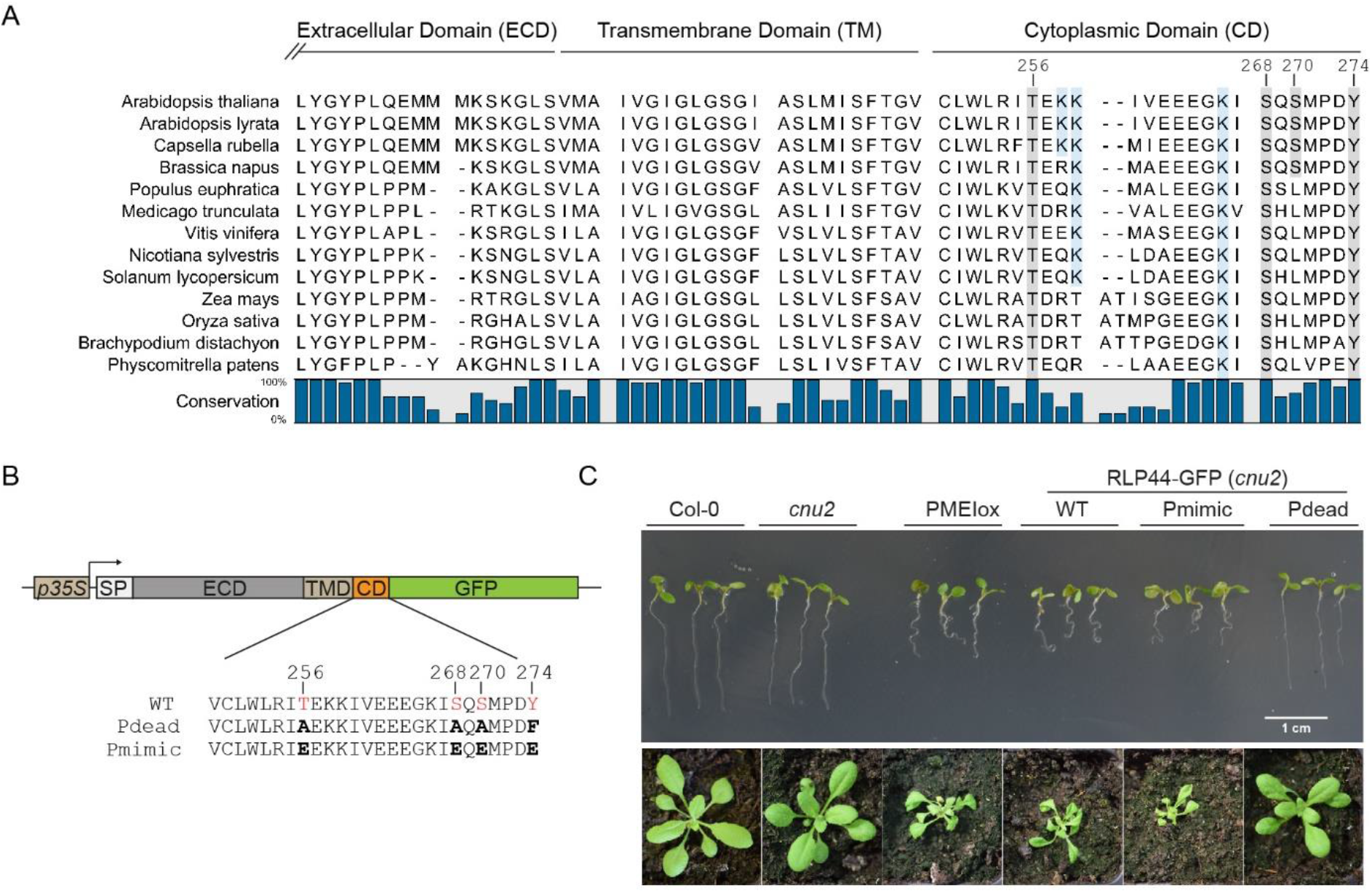
RLP44 has an unusual C-terminal tail with four putative phosphorylation sites that are required for function. **(A)**The short, cytoplasmic tail of AtRLP44 contains four putative phosphorylation sites, three of which are conserved in RLP44 orthologues. C-terminal part of the extracellular domain, predicted transmembrane domain (http://www.cbs.dtu.dk/services/TMHMM/), and cytoplasmic domain of AtRLP44 are indicated; see Supplemental Figure S1 for full alignment. **(B)** Schematic representation of RLP44 WT, Pdead, and Pmimic variants. **(C)** Blocking post translational modification of the four putative phosphosites in RLP44-GFP Pdead precludes function in the BR signalling-dependent response to cell wall modification, whereas simulating phosphorylation in RLP44-GFP Pmimic results in RLP44-GFP WT-like functionality. Expression of RLP44-GFP WT and Pmimic, but not of Pdead, is able to complement the PMEIox suppressor mutant *cnu2* and leads to recovery of the PMEIox root waving phenotype in seedlings and contorted leaf arrangement in adult plants.

**Figure 2.**
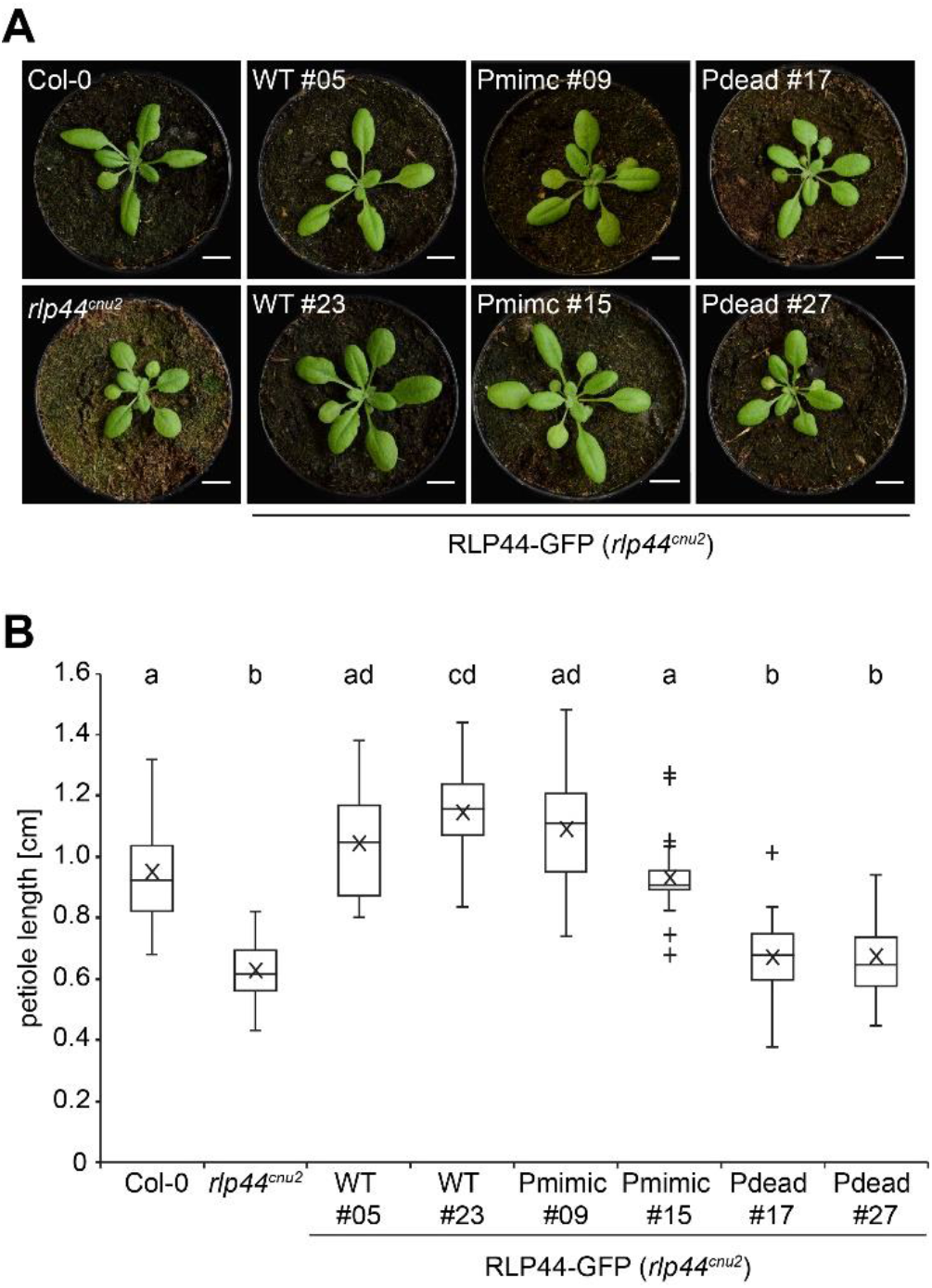
RLP44-GFP Pdead is unable to complement the petiole phenotype of *rlp44^cnu2^*. A) Rosette phenotype of Col-0, *rlp44^cnu2^*,and *rlp44^cnu2^* complementation lines in long day conditions. B) Quantification of petiole length in lines depicted in A). Boxes indicate range from 25^th^ to 75^th^ percentile, horizontal line indicates the median, whiskers indicate data points within 1.5 times the interquartile range. Markers above whiskers indicate outliers, n = 21. Lettering indicates statistically significant difference according to Tukey’s post-hoc test following one-way ANOVA.

### Phospho-status affects the subcellular localization of RLP44-GFP

We have previously shown that RLP44-GFP is localized in intracellular vesicles and at the plasma membrane (Wolf et al.,2014), in agreement with its association with receptors for extracellular signalling ligands. To assess whether modification of the four putative phosphorylation sites influences the subcellular localization of RLP44-GFP, we imaged root tips of transgenic lines expressing RLP44-GFP WT, Pdead or Pmimic in the wild-type (Col-0) background. RLP44-GFP WT showed the expected distribution of plasma membrane and intracellular fluorescence, which partially co-localized with the styryl dye FM4-64 (Figure 3A). Surprisingly, RLP44-GFP Pdead showed only very faint plasma membrane fluorescence and almost exclusive labelling of intracellular vesicles that appeared to largely co-localize with FM4-64 30 minutes after its application, suggesting endosomal localization (Figure 3A). In sharp contrast, RLP44-GFP Pmimic showed strongly enhanced plasma membrane labelling with few intracellular vesicles (Figure 3A). Similar results were observed in the *cnu2* background (Figure 3B) and confirmed by quantification of the mean plasma membrane to intracellular fluorescence ratio (Figure 3C).

**Figure 3.**
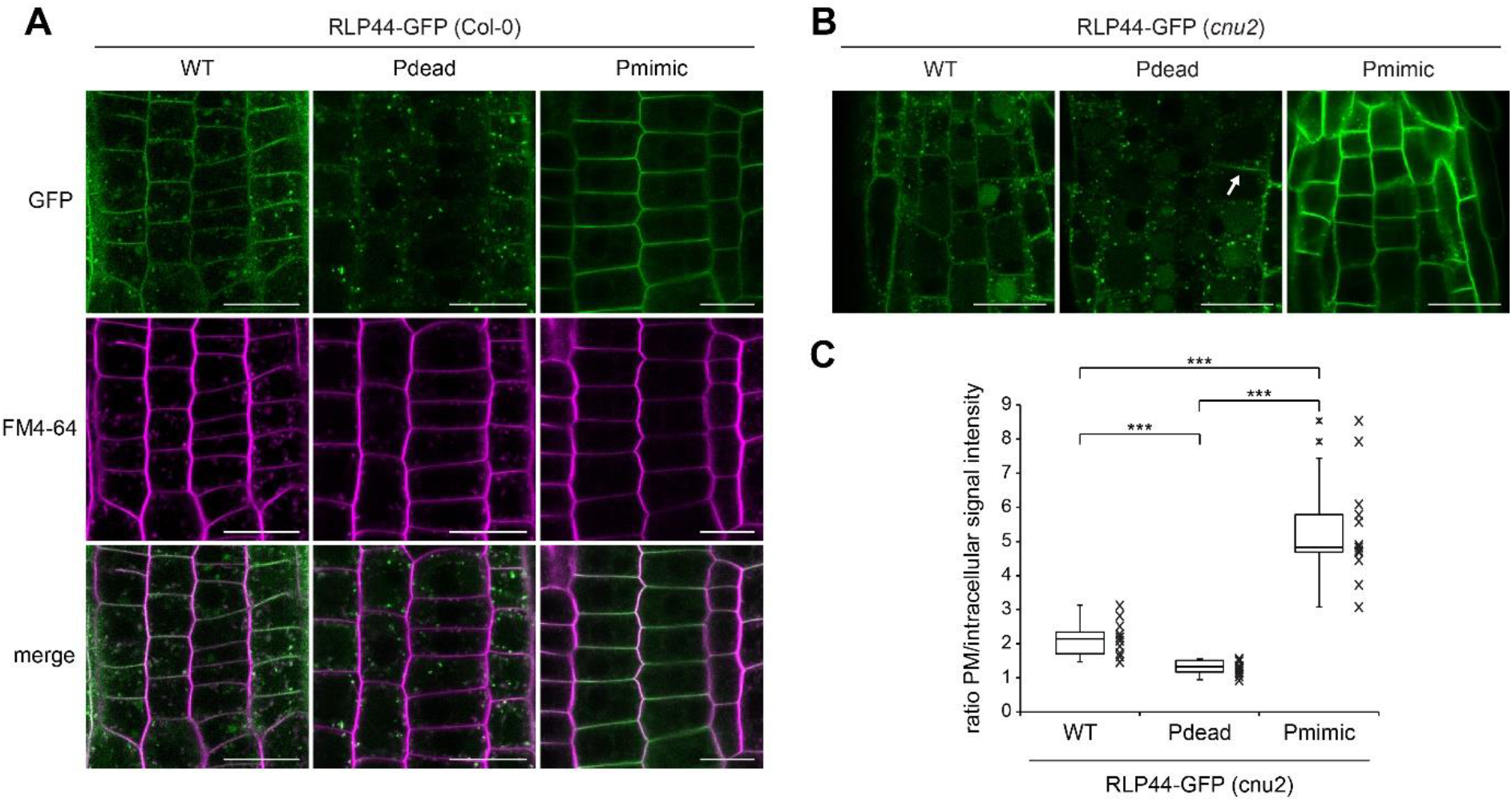
RLP44-GFP phospho-status determines its subcellular localization. (A) RLP44-GFP WT-derived fluorescence is observed at the plasma membrane and in the endomembrane system, as previously demonstrated (Wolf et al., 2014). RLP44-GFP Pdead is mostly confined to endomembranes, whereas RLP44-GFP Pmimic shows prominent plasma membrane localization. Bars = 20 μm. (B) The subcellular distribution of RLP44-GFP variants is maintained in the *cnu2* background. Arrow points to a fusing cell plate that shows increased RLP44-GFP Pdead fluorescence. Bars = 50 μm. (C) Quantification of mean plasma membrane to intracellular fluorescence ratio. Boxes indicate range from 25^th^ to 75^th^ percentile, horizontal line indicates the median, whiskers indicate data points within 1.5 times the interquartile range. Markers above whiskers indicate outliers, markers alongside box and whisker diagrams indicate individual data points, n = 12 measurements (cells) in 3 independent roots for each genotype. Asterisks indicate statistically significant differences with ***p<0.001 according to a Student’s t-test.

These results would be consistent with two different and mutually exclusive scenarios: first, the putative phosphorylation sites might be required for targeting of RLP44-GFP, thus the Pdead fluorescence distribution might reflect a failure of the receptor to be transported to the plasma membrane, whereas mimicking phosphorylation in Pmimic might enhance transport to the surface; second, differential rates of endocytic uptake might underlie the contrasting behaviour of the Pdead and Pmimic versions of RLP44-GFP. In the latter scenario, Pdead would be transported to the plasma membrane initially, but would rapidly undergo endocytosis, whereas mimicking phosphorylation would block internalization. As the second scenario is favoured by an increased Pdead fluorescence in expanding and fusing cell plates (Figure 3C), which exhibit rearrangement of cellular trafficking towards secretion (Richter et al., 2014), we sought to test the hypothesis that the three RLP44-GFP versions differ in their rate of endocytosis. To this end, we first interfered with endosomal trafficking by applying the phosphoinositide 3-kinase inhibitor Wortmannin (Wm). This treatment leads to a swelling of the multi-vesicular bodies (MVBs)/late endosomes (LE), thus indicating late endosomal nature of sensitive structures (Wang et al., 2009; Viotti et al., 2010). In addition, Wm can indirectly lead to an inhibition of endocytosis (Emans et al., 2002). All three RLP44-GFP versions were sensitive to Wm, as subpopulations of intracellular GFP-positive punctate showed pronounced swelling, suggesting that RLP44-GFP reaches late endosomes (Figure 4A). Moreover, plasma membrane labelling of RLP44-GFP Pdead was increased after Wm treatment. This is consistent with the hypothesis that this RLP44 version displays low steady state abundance at the plasma membrane because it experiences increased endocytic uptake. Consequently, inhibition of this uptake, in this case by Wm, leads to an increase in abundance at the plasma membrane (Figure 4A). To independently corroborate these results, we made use of the fungal toxin brefeldin A (BFA), which, in Arabidopsis roots, leads to aggregation of endosomal compartments into a hybrid organelle, the BFA compartment, in which endocytic cargo becomes trapped (Geldner et al., 2003; Grebe et al., 2003; Dettmer et al., 2006; Viotti et al., 2010). Consequently, quantification of fusion protein-derived fluorescence in BFA compartments has been used to assess endocytosis of plasma membrane receptors (Di Rubbo et al., 2013; Martins et al., 2015). After 120 minutes of BFA treatment, the three RLP44-GFP versions displayed differential accumulation in BFA compartments, with the Pdead version showing the strongest signal, followed by WT and Pmimic (Figure 4B, C). These observations are consistent with differential endocytosis as the mechanistic explanation for the different subcellular distribution of RLP44-GFP WT, Pdead, and Pmimic. In order to directly test the impact of retrograde trafficking on RLP44-GFP localization through genetic interference with endocytic uptake, we used a previously described line expressing, in an inducible manner, artificial miRNAs directed against the TPLATE adapter complex to block clathrin-mediated endocytosis (Van Damme et al., 2011; Gadeyne et al., 2014). As expected, induction of amiRNA expression for 48 hours led to a marked increase in the ratio of plasma membrane-localized to intracellular FM4-64 fluorescence and a dramatic reduction in size and quantity of FM4-64-positive BFA compartments (Figure 5A, B), indicating strongly decreased endocytosis. Importantly, amiRNA induction in plants expressing RLP44-GFP Pdead led to the appearance of GFP fluorescence at the plasma membrane, in sharp contrast to mock treatment (Figure 5A, B). Quantification of plasma membrane and intracellular GFP fluorescence in RLP44-GFP WT and Pdead lines revealed that inhibition of clathrin-mediated endocytosis leads to a WT-like fluorescence distribution of RLP44-GFP Pdead (Figure 5B), suggesting that phospho-status governs endosomal trafficking of RLP44-GFP.

**Figure 4.**
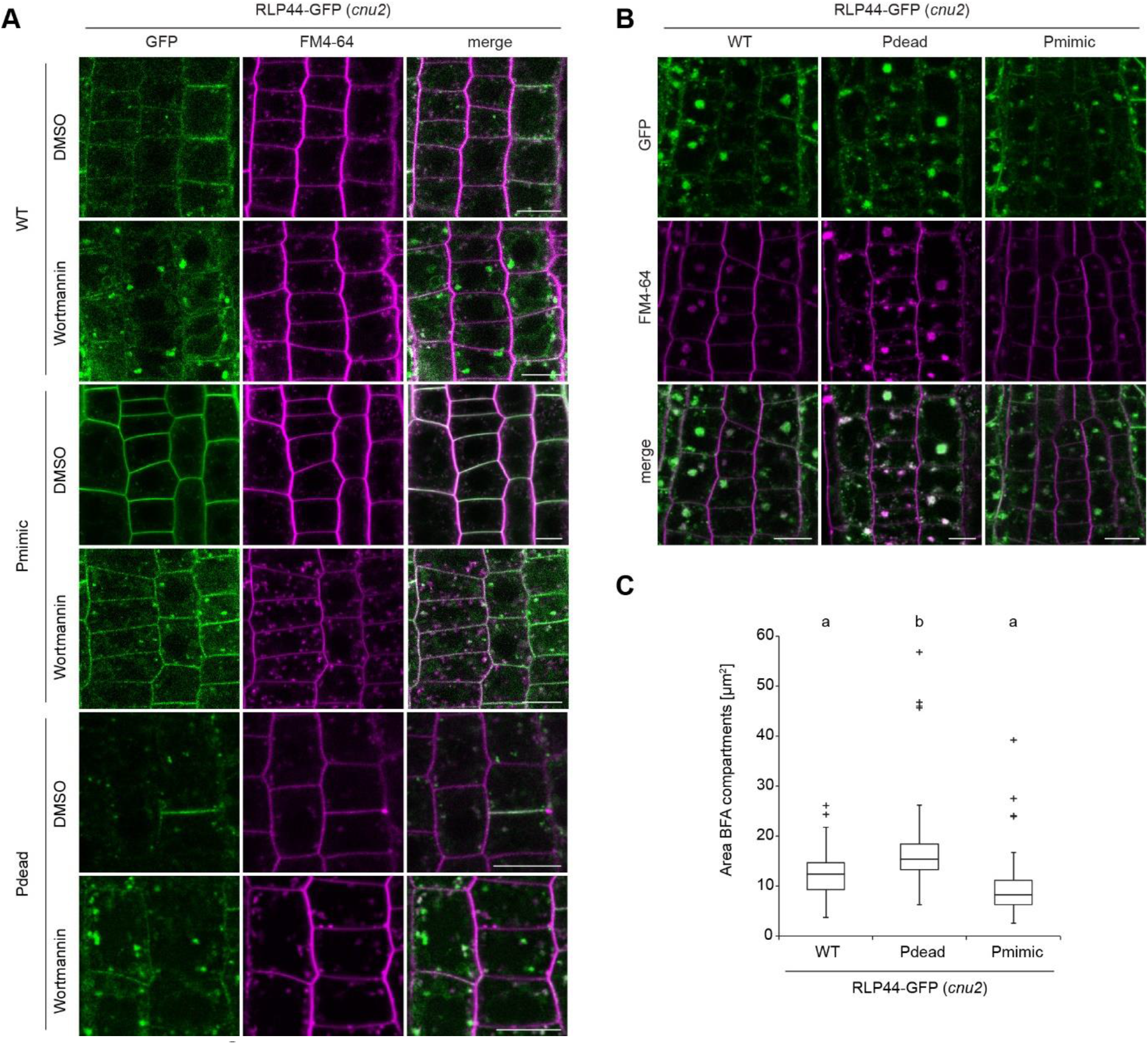
RLP44-GFP variants undergo endocytosis. **(A)** Fluorescence derived from RLP44-GFP WT, Pdead, and Pmimic variants accumulates in enlarged structures after 20 μM Wortmannin treatment for 165 minutes, suggesting they reach late endosomes. Note increased plasma membrane labelling of RLP44-GFP Pdead after WM treatment. Bars = 10 μm. **(B)** Fluorescence derived from RLP44-GFP WT, Pdead, and Pmimic variants accumulates in BFA bodies. Roots were treated with 50 μM of BFA or DMSO for 120 minutes and with FM4-64 for 20 minutes before imaging. **(C)** Image quantification reveals largest fluorescent area in BFA bodies of RLP44-GFP-derived fluorescence (lower panel), n = 90-116 measurements (cells) in 18 independent roots for each genotype. Lettering indicates statistically significant difference according to Mann-Whitney U-tests. Bars = 10 μm.

**Figure 5.**
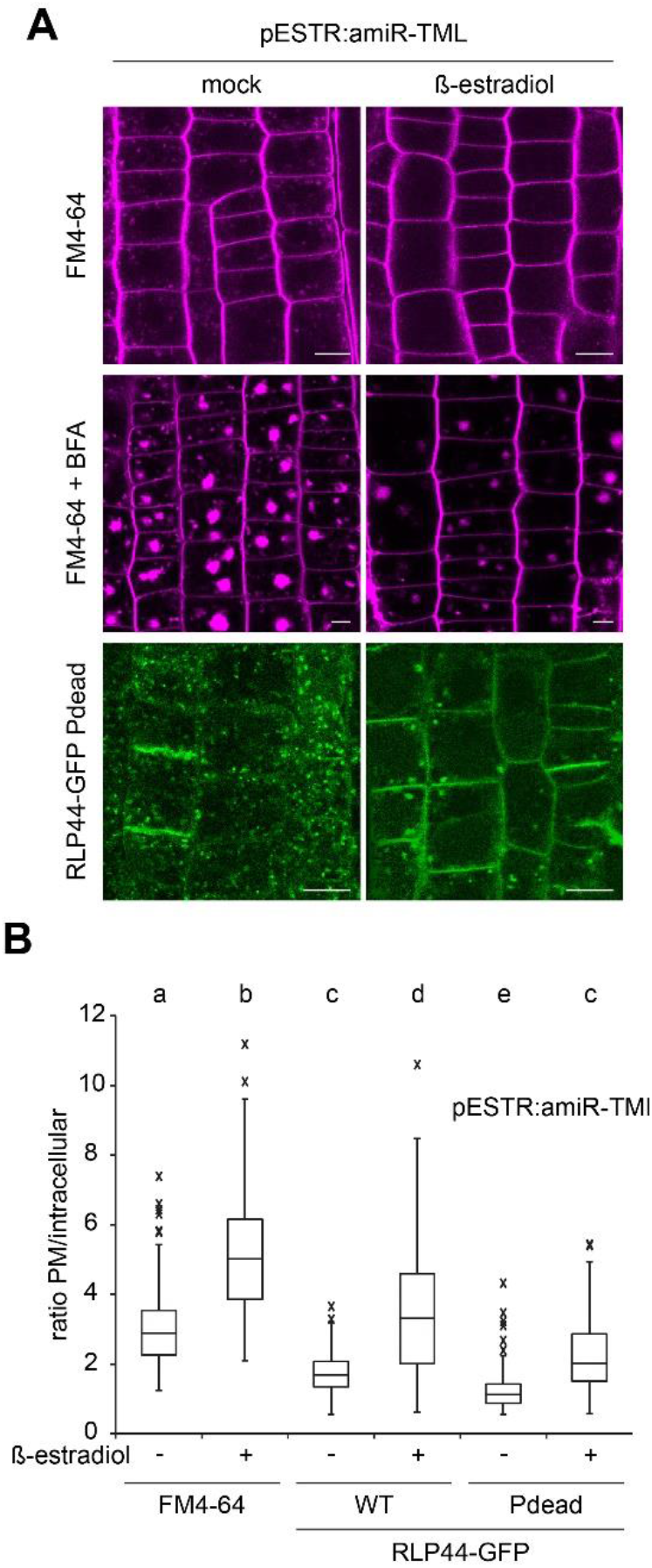
Blocking phosphorylation promotes endocytosis of RLP44-GFP. (A) Inducible expression of amiRNAs targeting the clathrin adapter complex TPLATE leads to reduced formation of endosomes and decreased uptake of FM4-64 into BFA bodies compared to mock control, suggesting efficient blockage of clathrin-mediated endocytosis. Knock-down of TPLATE in RLP44-GFP Pdead leads to enhanced plasma membrane localization. Bars = 10 μm. (B) Quantification of mean plasma membrane to intracellular fluorescence ratio in the indicated genotypes with and without amiRNA-mediated knock down of TPLATE. Boxes indicate range from 25^th^ to 75^th^ percentile, horizontal line indicates the median, whiskers indicate data points within 1.5 time the interquartile range. Markers above whiskers indicate outliers, n = 78-104 measurements from 15-18 independent roots each. Letters indicate statistically significant differences according to Dunn’s post hoc test after one-way Kruskal-Wallis test.

### Effect of phospho-status on membrane localization occurs independently of ubiquitination

Endocytic trafficking of LRR receptors often depends on mono or poly-ubiquitination (Lu et al., 2011; Martins et al., 2015; Zhou et al., 2018). We therefore tested whether RLP44 is ubiquitinated and whether phospho-status affects this protein modification. To this end, we immunopurified RLP44-GFP from transgenic plants and performed Western Blotting with antibodies against GFP and ubiquitin, respectively. Similar to other membrane-localized LRR proteins (Lu et al., 2011; Martins et al., 2015), RLP44 immunoprecipitates showed a ladder-like pattern of high molecular weight proteins detected in the anti-ubiquitin blot that are not present in the GFP-Lti6b control. (Figure 6A). To assess whether ubiquitination is affected by RLP44-GFP phospho-status we immunopurified the WT, Pdead, and Pmimic RLP44-GFP versions and compared the pattern and abundance of protein species reacting with the ubiquitin antiserum. Although the patterns differed slightly, all three variants clearly showed ubiquitinated forms in similar abundance, demonstrating that phosphorylation is neither required for, nor inhibiting ubiquitination (Figure 6B). Thus, the plasma membrane localization of RLP44-GFP Pmimic is not explained by an altered UB profile. As decoration with ubiquitin is not only involved in the initial internalization step of cell surface receptors, but also in the endosomal sorting and transport to the vacuole, the differing pattern of RLP44-GFP Pdead could be related to its largely endosomal localization.

**Figure 6.**
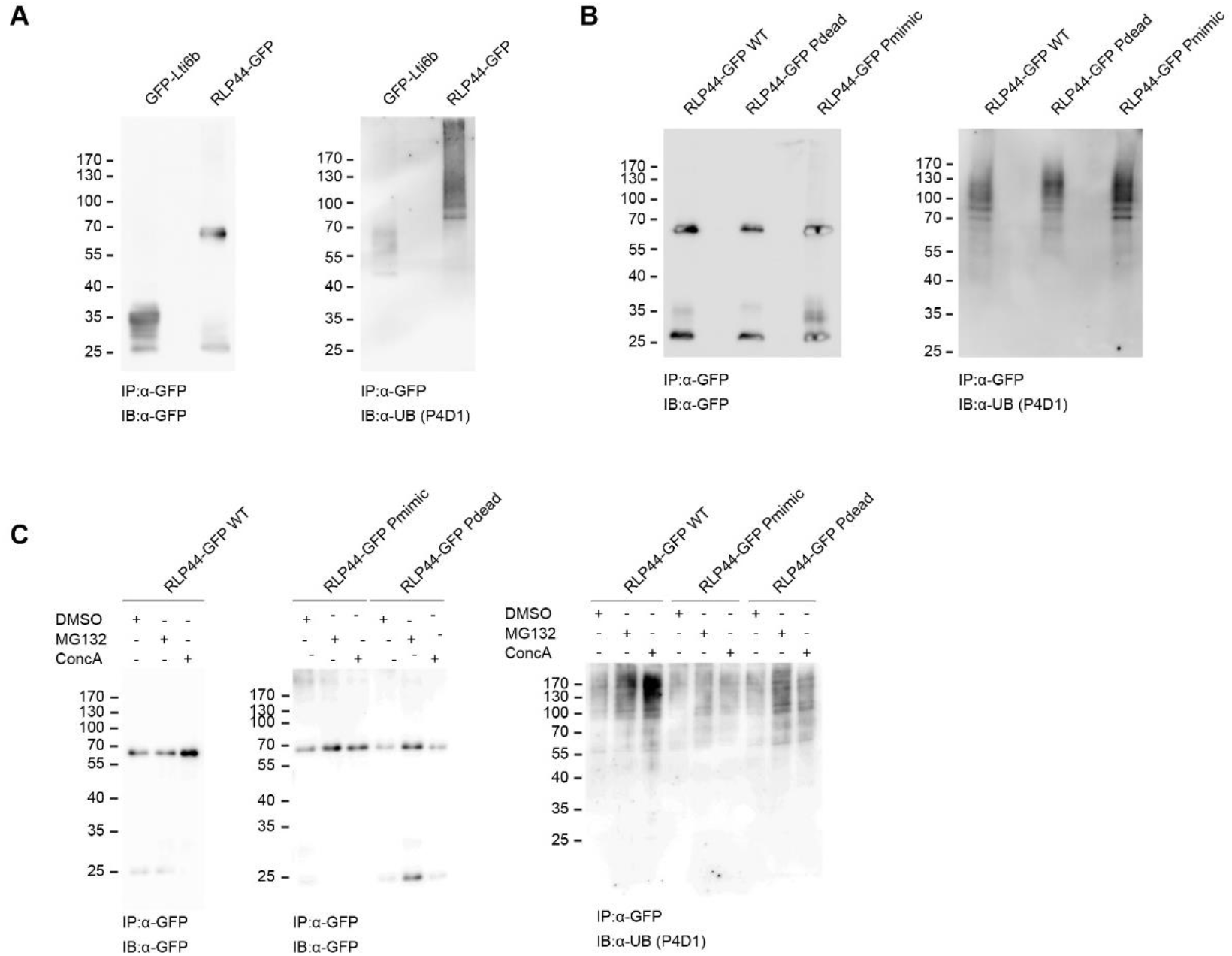
Phospostatus controls RLP44 localization independent of its decoration with ubiquitin. **(A)** Western Blotting of immunoprecipitates from GFP-Lti6b and RLP44-GFP seedlings probed with antibodies raised against GFP and ubiquitin, respectively, shows ubiquitin decoration of RLP44-GFP. **(B)** RLP44-GFP WT, Pdead ad Pmimic show similar patterns of ubiquitination. **(C)** Treatment with MG132 and ConcA increases the abundance of ubiquitinated protein species in immunoprecipitates of RLP44-GFP WT and RLP44-GFP Pdead, but not of RLP44-GFP Pmimic.

The proteasome inhibitor MG132 and the V-ATPase inhibitor ConcanamycinA (ConcA) are known to interfere with endosomal sorting and degradation of internalized receptors, increasing the abundance of UB-conjugated and thus presumably endosomal protein species (Dettmer et al., 2006; Lu et al., 2011; Luo et al., 2015; Zhou et al., 2018; Song et al., 2019). Consistent with this, treatment with MG132 or ConcA increased the abundance of ubiquitinated protein in immunopurified RLP44-GFP samples. Similar results were obtained with RLP44-GFP Pdead samples, in line with significant endosomal GFP fluorescence observed in these two lines. Conversely, the abundance of ubiquitinated protein did not increase after inhibitor treatment in the RLP44-GFP Pmimic line, which shows strongly reduced intracellular fluorescence (Figure 6C). In summary, phospho-status governs subcellular localization of RLP44 independent of ubiquitination.

### Subcellular distribution of RLP44 is modified by BRI1

To assess whether the subcellular localization of RLP44-GFP can be affected by the presence of its interaction partners, we quantified the subcellular distribution of RLP44-GFP fluorescence in the absence of BRI1. To this end, we crossed RLP44-GFP WT, Pdead, and Pmimic into the T-DNA insertion line *bri1-null* (Jaillais et al., 2011a), which, based on the available evidence, completely lacks BRI1 protein. Whereas neither localization of the RLP44-GFP WT version nor that of RLP44-GFP Pdead was strongly affected in the *bri1-null* mutant (Figure 7A, B), the almost exclusive plasma membrane localization of RLP44-GFP Pmimic clearly depended on the presence of BRI1 (Figure 7C): intracellular labelling was strongly increased in *bri1-null* compared to the Col-0 background (Figure 7A) and the plasma membrane-to-cytosol fluorescence ratio of Pmimic was not increased compared to the RLP44-GFP WT in the mutant background (Figure 7B). These results suggest that, at least in part, presence of BRI1 precludes endocytosis of the Pmimic version of RLP44. Accordingly, the behaviour of the Pdead variant, which is unable to promote BR signalling, is unaffected by the presence or absence of BRI1.

**Figure 7.**
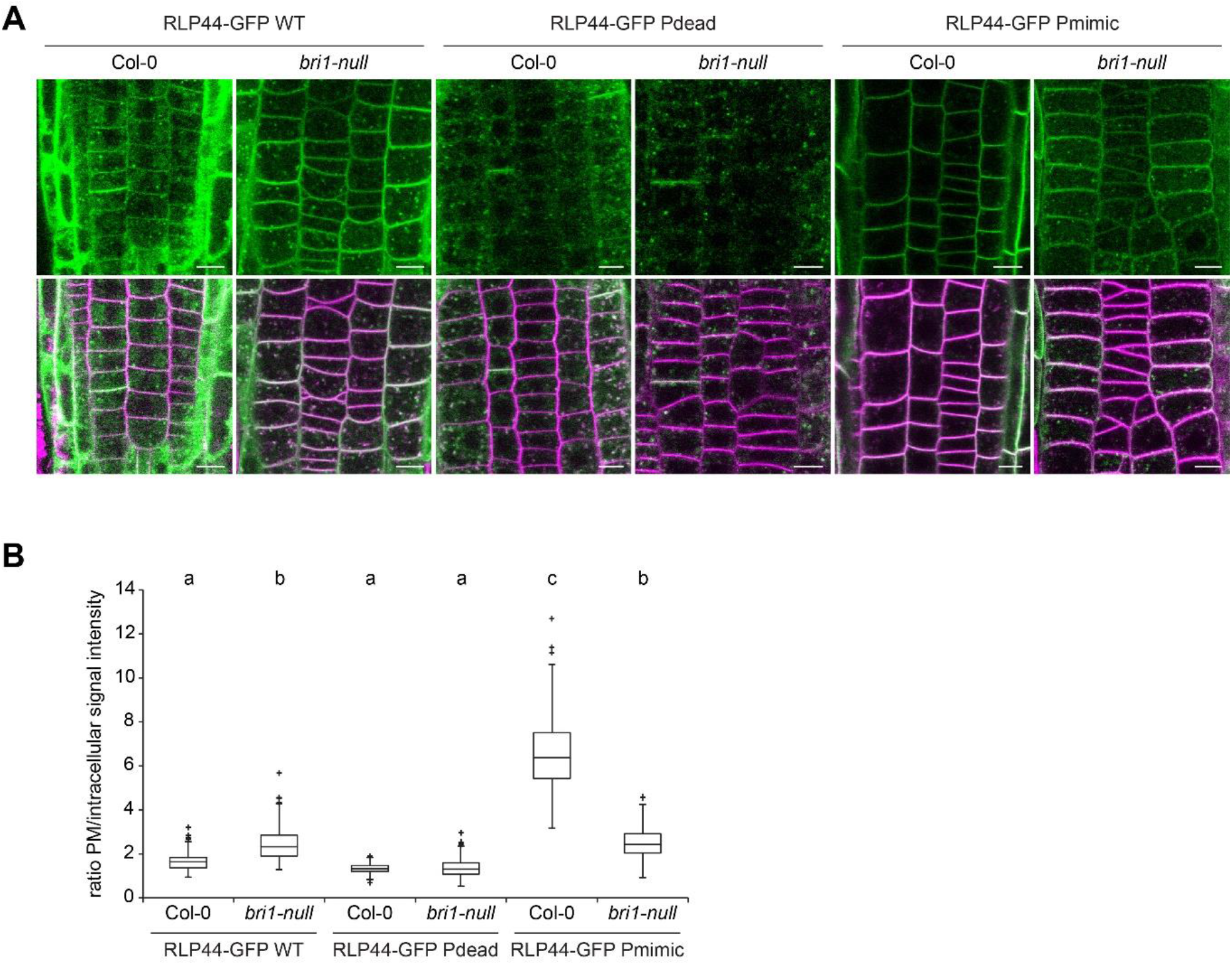
Subcellular distribution of RLP44-GFP depends on BRI1. **(A)** Comparison of fluorescence distribution derived from the three RLP44-GFP variants in Col-0 and *bri1-null* background. Note the increased appearance of intracellular fluorescence in RLP44-GFP Pmimic expressed in the *bri1-null* mutant. Bars = 5 μm. **(B)** Quantification of mean plasma membrane to intracellular fluorescence ratio in the indicated genotypes, derived from images as in A). Boxes indicate range from 25^th^ to 75^th^ percentile, horizontal line indicates the median, whiskers indicate data points within 1.5 time the interquartile range. Markers above whiskers indicate outliers, n = 48-62 cells from 12 independent roots for each genotype. Letters indicate statistically significant differences according to Tukey’s post hoc test after one-way ANOVA.

### Phosphorylation is not required for the role of RLP44 in PSK signalling

We have recently described (Holzwart et al., 2018) that, apart from promoting BR signalling, RLP44 plays a role in promoting signalling mediated by the receptors of PSK peptide hormones, PSKR1 and PSKR2 (Matsubayashi et al., 2002; Matsubayashi et al., 2006). Having demonstrated that the role of RLP44 in BR signalling depends on the four phosphosites in the cytosolic tail, we next assessed whether the Pdead and Pmimic versions affected functionality of RLP44 in the PSK pathway. To this end, we analysed PSK-related phenotypes in complemented *rlp44^cnu2^* mutants. PSK signalling is required in the epidermis for normal root elongation (Kutschmar et al., 2009; Hartmann et al., 2013) and exogenously applied PSK peptide leads to a moderate increase in the root length of wild type seedlings (Kutschmar et al., 2009; Ladwig et al., 2015; Wang et al., 2015). In line with its role in promoting PSK signalling, *rlp44^cnu2^* is compromised in this response (Holzwart et al., 2018; Holzwart et al., 2020). Analysis of the complementation lines described earlier (Figure 2) revealed that all three RLP44-GFP versions, including Pdead, were able to restore the response to PSK in *rlp44^cnu2^* (Figure 8A), suggesting that PSK and BR signalling might have different requirements for the modification of the four RLP44 phosphorylation sites. Notably, the PSK signalling-mediated promotion of root growth is believed to occur in the epidermis, a tissue where *RLP44, BRI1* and *PSKR1* are co-expressed (Friedrichsen et al., 2000; Matsubayashi et al., 2006; Kutschmar et al., 2009; Holzwart et al., 2018). To corroborate these results, we tested whether RLP44-GFP Pdead could complement the ectopic xylem phenotype of *rlp44^cnu2^* seedling roots, which is caused by reduced PSK signalling (Holzwart et al., 2018). In line with the root length assay, expression of RLP44-GFP Pdead led to wild type-like xylem cell numbers (Figure 8B), confirming that phosphorylation of these residues is not a requirement for the role of RLP44 in promoting PSK signalling. To exclude a PSK signalling-independent effect of RLP44 Pdead on xylem cell numbers we performed a rescue experiment in *pskr1-3 pskr2-1* double mutants. As previously described for the WT RLP44-GFP (Holzwart et al., 2018), RLP44 Pdead requires the presence of PSK receptors to exert any effect on xylem cell number (Figure 8C). In summary, our results suggest that phosphorylation modulates RLP44 functionality in different receptor complexes.

**Figure 8.**
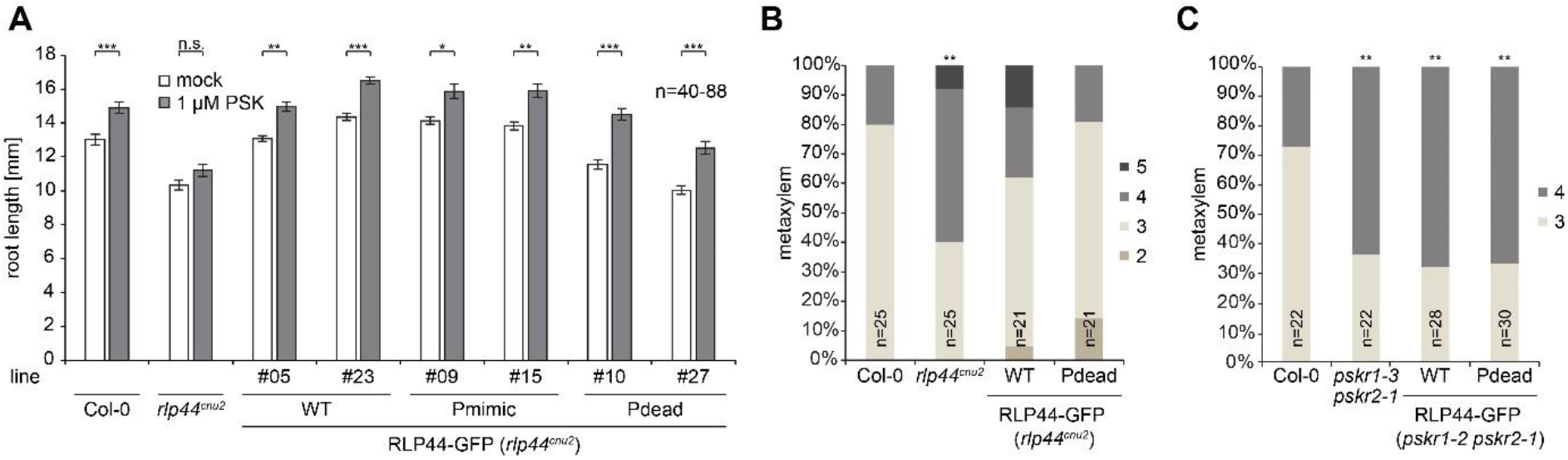
RLP44 is functional in PSK signalling irrespective of potential phospho-status. **(A)** Root length response to exogenous application of PSK peptide is impaired in *rlp44^cnu2^* (Holzwart et al., 2018), but restored by expression of all three RLP44-GFP variants, including Pdead. Bars denote root length after five days of growth on control plates or plates containing 1 μM PSK peptide, n = 40-88. Asterisks indicate statistical significance according to Tukey’s post hoc test after one-way ANOVA with *P<0.05, **P<0.01, ***P<0.001. **(B)** The PSK signalling-dependent ectopic xylem phenotype of *rlp44^cnu2^* (Holzwart et al., 2018) is restored by expression of RLP44-GFP Pdead. **(C)** Effect of RLP44 expression depends on the presence of PSK receptors PSKR1 and PSKR2. Bars in B) and C) denote frequency of roots with indicated number of metaxylem cells, asterisks indicate statistically significant difference from Col-0 based on Dunn’s post-hoc test with Benjamini-Hochberg correction after Kruskal-Wallis modified U-test (**p < 0.01).

### Phospho-charge, rather than modification of individual amino acids determines RLP44 functionality in BR signalling

During the course of this work, we noticed that a previously characterized RLP44 line (Holzwart et al., 2018, 2020) in which a serine-rich linker ((GS)_11_) separates RLP44 and GFP, tended to show strong overexpression phenotypes despite being under control of the native RLP44 promoter and localized predominantly at the plasma membrane (Supplemental Figure S5A, B). To test whether serine residues in this linker might become phosphorylated, we performed phosphatase assays on immunoprecipitated RLP44 protein fusions. In contrast to RLP44-GFP (Supplemental Figure S3B), RLP44-(GS)_11_-GFP harbouring the serine-rich linker clearly showed an additional band of lower mobility, which disappeared after calf intestinal phosphatase (CIP) treatment, indicating that the upper band represents a phosphorylated form of the fusion protein (Figure 9A). These results suggest that the immediate environment of the RLP44 C-terminus can undergo phosphorylation and that hyperphosphorylation in the linker might confer a gain-of-function phenotype with respect to BR signalling. Moreover, a hyperphosphorylated linker can be exploited to visualize regulation of phosphorylation. Western blotting after immunoprecipitation revealed that the relative abundance of the lower mobility (i.e. phosphorylated) form of RLP44-(GS)_11_-GFP seemed to be reduced by treatment with the kinase inhibitor K252a and increased by treatment with brassinolide (Figure 9B). To further elucidate how hyperphosphorylation of RLP44 affects activity, we created Pdead and Pmimic versions of RLP44-GFP harbouring the serine-rich linker. In contrast to what was observed with the previous RLP44-GFP constructs, all three RLP44-(GS)_11_-GFP version complemented the *cnu2* rosette phenotype and showed almost exclusive plasma membrane localization (Figure 8D, E). This suggests that hyperphosphorylation increases RLP44 activity with respect to BR signalling and shifts its localization to the plasma membrane. In addition, these results would suggest that overall charge, rather than modification of specific amino acids, are the determining factor in controlling RLP44 activity.

**Figure 9.**
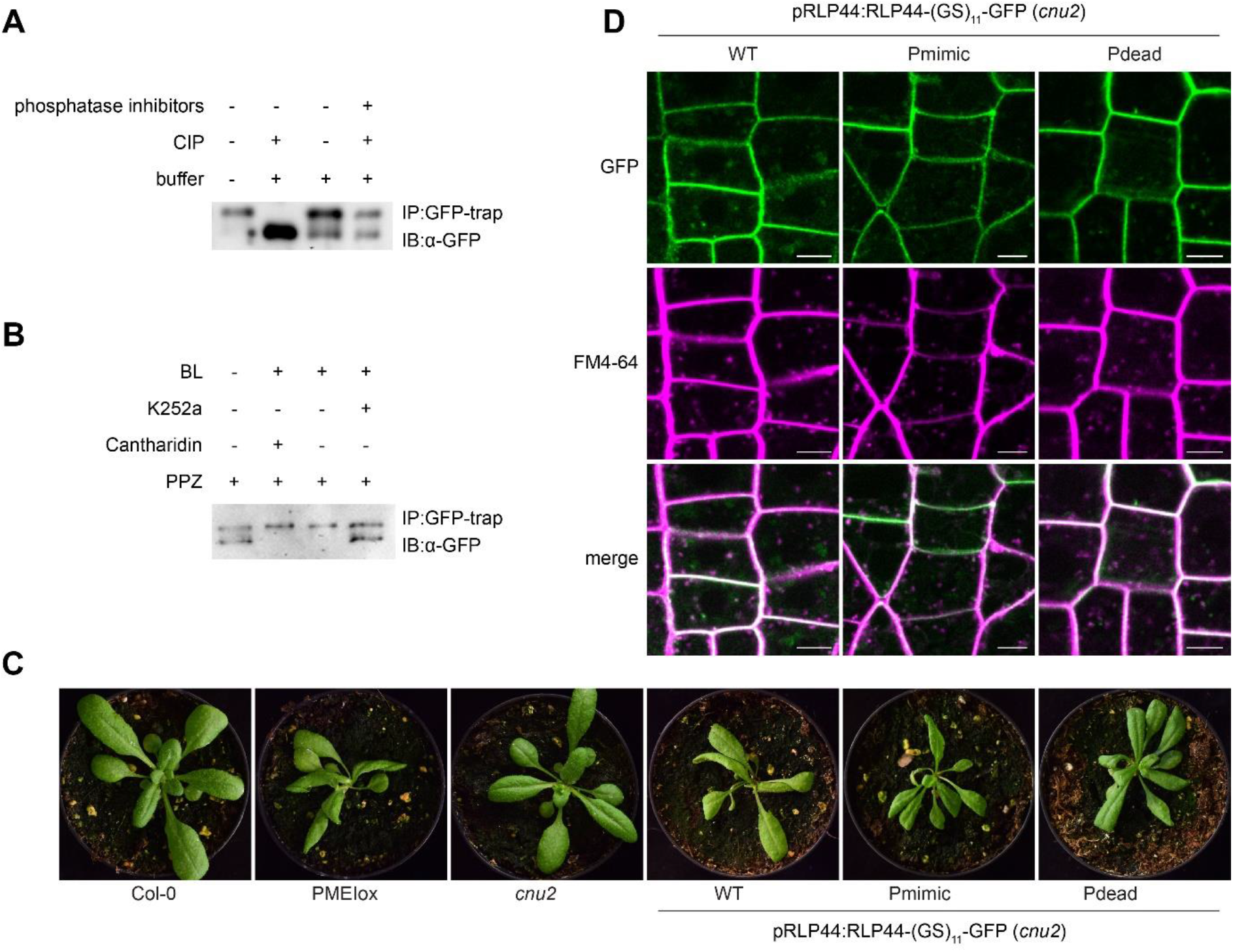
Overall charge, rather than specific phosphosites modulate RLP44 function. **(A)** Phosphatase treatment of pRLP44:RLP44-(GS)_11_-GFP leads to a pronounced shift in electrophoretic mobility, visualized by western blotting using GFP antiserum. **(B)** The relative abundance of the lower mobility, phosphatase-sensitive RLP44-(GS)_11_-GFP band is responsive to BL and kinase inhibitor treatment. **(C)** The Pdead version of pRLP44:RLP44-(GS)_11_-GFP is able to complement *cnu2* in a similar manner as the WT and Pmimic versions. **(D)** All three RLP44-(GS)_11_-GFP versions mainly accumulate at the plasma membrane.

It is not surprising that attaching a C-terminal fusion partner to a receptor protein might alter its activity in different ways. Therefore, the three 35S-driven RLP44-GFP versions might accentuate phosphorylation-mediated tuning of RLP44 activity at the plasma membrane. To elucidate the role of RLP44 phosphorylation independent of the linker sequence, we generated constructs encoding untagged RLP44 with single mutations in individual phosphosites as well as quadruple Pmimic and Pdead versions. We used these constructs along with the WT RLP44 version to complement *cnu2* (Wolf et al., 2014) and scored the aberrant cellular morphology phenotype described for PMEIox (Li et al., 2021), as this represents the most sensitive read-out of RLP44 function. As expected, we found that the RLP44 WT construct led to a reversion of the *cnu2* phenotype, resulting in root meristems that were indistinguishable from PMEIox (Figure 10). In contrast, RLP44 versions in which individual phosphosites were mutated to alanine (T256A, S268A, A270A) or phenylalanine (Y274F) resulted in strongly reduced reversion of the *cnu2* phenotype (Figure 10). A similar phenotype was displayed by the Pdead version, while Pmimic fully complemented *cnu2*. Even though these experiments suggest a more nuanced effect of the phosphosites in untagged RLP44, they broadly confirm our previous results. It should also be noted that, based on QRT-PCR analysis, all untagged complementation lines show rather high expression of the RLP44 transgene, despite being driven from the *RLP44* promoter (Supplemental Figure S5C). There are a several conceivable explanations for this, ranging from an incomplete, hyperactive promoter fragment to increased mRNA stability in the transgene due to the recombinant 3’ environment. Nevertheless, these results are consistent with the notion that RLP44 function is modified by the phospho-state of its C-terminus. Taken together, our results show that availability of RLP44 to engage with the BRI1 or PSKR1 receptor complexes is differentially modulated by phosphorylation and demonstrates how selectivity and specificity of different receptor-mediated signalling pathways can be encoded in the pattern of post-translational modifications.

**Figure 10.**
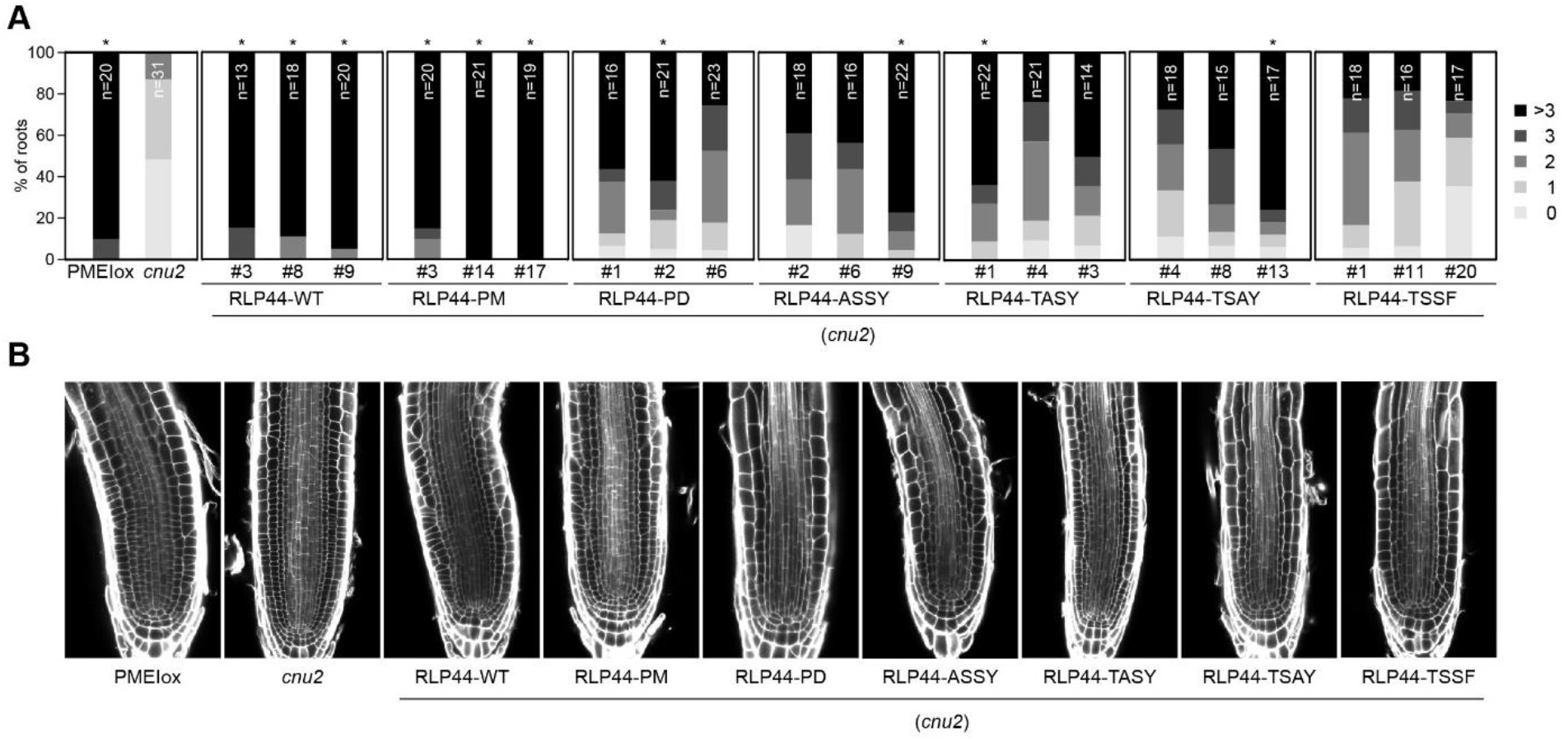
RLP44 phosphosites are crucial for its function in the native protein context. **(A)** Mutation of individual putative phosphosites in untagged RLP44 constructs results in reduced restoration of the PMEIox cell wall orientation phenotype in the *cnu2* background. Upper panels show quantification of aberrant cell wall orientation in median sections of root meristems. Asterisks indicate statistically significant difference from *cnu2* according to Kruskal-Wallis non-parametric test followed by Chi-square analysis after Bonferroni adjustment **(B)** Exemplary root meristem CLSM images of the indicated genotypes after staining with Calcofluor White and ClearSee (Kurihara et al., 2015) treatment.

## Discussion

Plants rely on a complex network of cell surface receptors to integrate developmental and environmental cues into behaviour adapted to the conditions. In light of the many possible interactions between LRR-RLKs (Smakowska-Luzan et al., 2018) and the high likelihood of accidental encounters in the crowded plasma membrane, which restricts mobility to two dimensions, a central question in signal transduction is thus how a specific response that transcends noise is ensured. In the current model of LRR-RLK signalling activation, the extracellular ligands serve as molecular tether to bring receptor and co-receptor together and thereby juxtapose their kinase domains in the cytosol, so that trans-phosphorylation can ensue (Hohmann et al., 2017). Fully activated kinase domains then recruit downstream signal transduction components which can be dedicated to individual signalling pathways (He et al., 2000; Brutus et al., 2010; Hohmann et al., 2018). For example, a chimeric receptor constituted by the BRI1 extracellular domain and the cytosolic domain of the immune receptor XA21 triggered immune signalling in response to BRs (He et al., 2000), whereas a similar BRI1-HAESA chimera was able to complement *haesa* mutants in a BR dependent manner (Hohmann et al., 2018). However, several pathways share components, as demonstrated by the ubiquity of SERK co-receptors (Ma et al., 2016), which are also an essential component of both PSK and BR signalling (Ladwig et al., 2015; Wang et al., 2015), the two LRR-RLK-governed pathways studied here. In addition to RLKs, RLPs contribute to receptor-mediated signalling and increase the complexity of the plasma membrane receptor network. We have previously demonstrated that RLP44 can promote both PSKR1 and BRI1-mediated signalling, but it was unclear how these activities are coordinated in light of the fact that RLP44 seems to act independently of the BR and PSK ligands (Wolf et al., 2014; Holzwart et al., 2018). Here, we show that phospho-status can route RLP44s towards functioning in PSK or BR signalling. This is consistent with previous proteomic analysis that revealed that putative phosphorylation sites are enriched in the binding interfaces of heterooligomers (Nishi et al., 2011) and proteins that engage in multiple (mutually exclusive) interactions using the same contact surface, presumable acting as a switch between different pathways (Tyagi et al., 2009). While we could not unequivocally demonstrate phosphorylation of T256, S270, and Y274, their modification by a negative charge generally seems to promote function in the BRI1-mediated signalling pathway.

Post-translational modifications play a central role in regulating the trafficking of plasma membrane proteins. It is well established that ubiquitination can act as a signal for internalization and endosomal sorting across kingdoms (Haglund and Dikic, 2012; Dubeaux et al., 2015). The 76 amino acid protein ubiquitin is linked via its C-terminal glycine to lysine residues in the target protein. Endosomal trafficking of the LRR-RLKs FLS2 and BRI1 has been shown to depend on ubiquitination (Lu et al., 2011; Martins et al., 2015; Zhou et al., 2018). Several prominent examples from the animal literature demonstrate the versatility of phosphorylation for the regulation of trafficking (Offringa and Huang, 2013). For instance, phosphorylation of epidermal growth factor receptor (EGFR) triggers interaction with the AP-2 adaptor complex and a E3 ubiquitin ligase, respectively, to promote endocytosis through two different pathways (Bakker et al., 2017). In plants, only for very few plasma membrane receptor proteins a direct link between phosphorylation and an effect on trafficking has been demonstrated. Endocytosis of LYSIN MOTIF-CONTAINING RECEPTOR-LIKE KINASE5 (LYK5) appears to be regulated by CHITIN ELICITOR RECEPTOR KINASE 1 (CERK1)-mediated phosphorylation (Erwig et al., 2017). In addition, a phospho-dead mutant of FLS2 (T867A) shows reduced endocytosis, and the same effect is observed when kinase activity is pharmacologically inhibited (Robatzek et al., 2006); however, ubiquitination is not dependent on phosphorylation of this residue (Lu et al., 2011). Besides FLS2, many LRR-RLKs undergo ligand binding-induced endocytosis. While a direct regulatory role of specific phosphorylation events, for example through mediating interaction with the ubiquitination machinery, has yet to be shown, receptor complex auto- and trans-phosphorylation is the primary output of ligand perception and thus likely to be involved in coupling receptor activation to endocytosis. Interestingly, RLP44 shows a highly conserved lysine at position 266. It is thus tempting to speculate that phosphorylation might affect ubiquitination (Swaney et al., 2013; Venne et al., 2014), as previously observed with plasma membrane proteins such as EGFR (Sigismund et al., 2013). However, the effect of phospho status on RLP44 membrane localization seems to occur independently of ubiquitin, as the plasma-membrane-localized RLP44-GFP Pmimic showed a ubiquitination pattern similar to that of the WT version. In addition, in our mass spectrometry data, we found the characteristic ubiquitin di-glycine remnant (after trypsin digestion) on Lys266 of both RLP44-GFP WT and RLP44-GFP Pdead. Although these data do not rule out an impact of phosphorylation on the type of ubiquitination, they do show that phosphorylation is not required for the decoration with ubiquitin *per se*, nor does the introduction of phosphorylation-like charge inhibit ubiquitination. Increasing the charge of the RLP44 C-terminus could promote plasma membrane localization through different means, favouring the interaction with BRI1. However, our results point to a slightly different scenario: RLP44 might be constitutively primed for endocytosis, and its dwell time at the membrane might be determined by the interactions it engages in. In favour of this hypothesis, endosomal uptake of RLP44-GFP Pmimic was strongly enhanced in the absence of BRI1. Notably, RLP44-GFP phospho-status had no impact on its functionality in the PSK pathway, despite RLP44-GFP Pdead showing very low abundance at the plasma membrane. This raises the question of whether PSKR1 signalling can occur from endosomes, as it is assumed for diverse signalling pathways in animals (Howe and Mobley, 2004; Sigismund et al., 2012) and also discussed in plants (Geldner and Jurgens, 2006). Alternatively, brief presence of RLP44-GFP Pdead at the plasma membrane might be sufficient for supporting PSK signalling. To assess whether membrane sub-compartmentalization contributes to the maintenance of specificity, it will be important to track trafficking of RLP44 together with its interaction partners BRI1 and PSKR1 using advanced imaging technology (Bucherl et al., 2017; McKenna et al., 2019). We found apparent orthologues of RLP44 in all analysed plant species, suggesting that RLP44-like genes are under strong selective pressure, similar to other RLPs involved in development such as CLV2 and TMM (Fritz-Laylin et al., 2005). Notably, T256, S268, Y274 are strictly conserved in all analysed RLP44 orthologues, despite the tendency of phosphorylation sites to diverge rapidly in linear motifs and receptor proteins (Holt et al., 2009; Riano-Pachon et al., 2010; Beltrao et al., 2012). In contrast, S270 is only found in *Brassicacea* and does not substitute an acidic residue, which is assumed to contribute a dynamic switch to a pre-existing interaction (Pearlman et al., 2011). In this respect, it is noteworthy that we identified RLP44 as required for the response to changes in pectin de-methylesterification in Arabidopsis (Wolf et al., 2014). This cell wall modification apparently evolved from an ancient cell wall consolidation mechanism and is operative in extant members of the charophytes (Proseus and Boyer, 2006; Popper et al., 2011; Wolf et al., 2012a; Domozych et al., 2014; Nishiyama et al., 2018). On the other hand, BRI1-like brassinosteroid receptors and PSK perception seem to be restricted to seed plants (Cheon et al., 2013) (Bowman et al., 2017), thus RLP44 orthologues could predate some of its interaction partners. It will be interesting to dissect how a protein like RLP44, which modulates the function of distinct RLKs, co-evolved with its interaction partners.

## Methods

### Plant material and growth conditions

All plants used in this study were of the Col-0 ecotype and are described in Table S1. Seeds were sterilized with 1.2% NaOCl in 70% ethanol and washed twice with absolute ethanol, and dried under sterile conditions. If not indicated otherwise, plants were grown in ½ strength MS medium supplemented with 1% sucrose and 0.9% plant agar. For treatment with ConcA and MG132, plants were grown for 6 days on standard medium on plates and transferred to liquid half-strnegth MS medium containing DMSO (mock treatment), 50 μM MG132 (Sigma), or 1 μM ConcA for 5 hours before sample collection and immunopurification. To study the seedling growth response to PSK, plates were supplied with 10 nM α-PSK (PolyPeptide). Roots were measured after 6 days of growth in long day conditions using Image J (https://imagej.nih.gov/ij/). Average or average normalized to the mock control together with the standard deviations of, at least 50 plants, were plotted for each experiment. To study the phosphorylation status of RLP44 in different mutant backgrounds, around 100 seeds per well were disposed in 6-well-plates (Corning) containing 5 mL of liquid ½ MS medium supplied with 1% sucrose. After stratification, plates were placed into growth chambers with long day conditions and kept in agitation (100 rpm). 3-days-after germination, seedlings were treated with 5 μM PPZ (Sigma-Aldrich). 6-days-after germination and 3-hours-before harvesting, seedlings were treated independently with 50 μM Cantharidin (Sigma-Aldrich), 2 μM K-252a (Enzo Life Sciences) or 50 μM of DMSO. 1.5-hours-before harvesting, seedlings were treated with 1 μM BL or 80% Ethanol. Afterwards, plant material was carefully collected, weight and directly froze in liquid N_2_.

### Cloning

RLP44-GFP under control of the CaMV 35S promoter was generated by amplifying the coding sequence of the (intronless) *RLP44* from genomic DNA using primers SW660 and SW670, and subsequent Gateway cloning into pDONR207 and pK7FWG2 (Karimi et al., 2002). RLP44-GFP Pdead was generated by introducing the T254A mutation with site-directed mutagenesis using primers SW666 and SW667 and RLP44 in pDONR207 as template. After recombination reaction into pK7FWG2, this plasmid was used as a PCR template to introduce the remaining three mutations with primers SW668 and SW660. The resulting PCR product was introduced in pDNOR207 through BP reaction, after which a sequence-confirmed clone was used for LR reaction into pK7FWG2. RLP44-GFP Pmimic was created analogously using primers SW672 and SW673 to introduce the T256E mutation and primers SW671 and SW660 to introduce the remaining mutations via PCR. The resulting PCR product was introduced in pDNOR207 through BP reaction, after which a sequence-confirmed clone was used for LR reaction into pK7FWG2. The pRLP44:RLP44-(GS)_11_-GFP Pdead and Pmimic constructs were generated by using primer SW1179 and SW1367 (Pdead) or SW1368 (Pmimic) with the Pdead and Pmimic constructs described above. The resulting PCR products were ligated into pGGC000 to generate entry clones. Destination clones for plant transformation were assembled with GreenGate cloning as described in Table S3. The WT pRLP44:RLP44-(GS)_11_-GFP construct was described before (Holzwart et al., 2018). In-fusion cloning with primers 2446-2455 was used to generate a second set of mutant constructs in entry vector pGGC000 (Lampropoulos et al., 2013), fusing RLP44 variants to GFP, separated by a GA-linker. The GFP portion was amplified with primers SW using vector pK7FWG2 as template. RLP44 fragments were amplified using the corresponding construct described above as template as follows: WT with SW 2446+2447, Pdead with SW2446 + 2454, Pmimic with SW2446 + SW2455, ASSY with SW2446 + SW2447, TASY with SW2446 + SW2447, TSAY with SW2446 + SW2450, TSSF with SW2446 + SW2452. GreenGate cloning was performed to generate RLP44 version under control of the RLP44 own promoter (Table S3). Untagged RLP44-WT, RLP44 Pdead, RLP44 Pmimic and RLP44 single mutations under control of the endogenous RLP44 promoter were generated by using the previously constructs as template for PCR reactions using primer SW3000 and one of the primers SW3001-3006 (Table S2). Subsequently, Gateway cloning into pDONR207 and pGWB501 was performed.

### Immunopurification and mass spectroscopy analysis

Immunopurification from Arabidopis seedlings and Western Blotting were performed as described previously (Holzwart et al., 2018), anti-ubiquitin antiserum (P4D1, Santa Cruz Biotechnology) was diluted 1:2000 in 1xTBST with 1% skim milk powder. For analysis of phosphorylation of the pRLP44:RLP44-(GS)_11_-GFP line, seedlings were grown in liquid culture for four days, harvested, and proteins were extracted das previously described (Holzwart et al., 2018). Treatment with calf intestinal phosphatase in the presence and absence of phosphatase inhibitors was carried out for one hour, after which samples were boiled in SDS sample buffer and subjected to Western Blotting as previously described (Holzwart et al., 2018).

For mass spectroscopy experiments, RLP44-GFP (WT or mutant variants) were transiently expressed in three-to four-week-old *N. benthamiana* leaves by agroinfiltration as previously described. Samples were taken two days post-infiltration; accumulation of the GFP-fused proteins was confirmed by confocal microscopy. Plant tissue was ground in liquid nitrogen, total proteins were extracted by adding lysis buffer (100 mM Tris-HCl pH 8.0; 150 mM NaCl; 10% glycerol; 5 mM EDTA; 5mM DTT, 1mM PMSF; 1% protease inhibitor cocktail; 1% NP-40) and the extracts were cleaned by filtration; extracts were incubated with GFP-Trap beads (Chromotek, Germany) for one hour, and beads were subsequently washed using washing buffer with detergent (100 mM Tris-HCL pH 8.0; 150 mM NaCl; 10% glycerol; 2mM DTT; 1% protease inhibitor cocktail; 0.2%NP-40) three times and washing buffer without detergent (100 mM Tris-HCL pH 8.0; 150 mM NaCl; 10% glycerol; 2mM DTT; 1% protease inhibitor cocktail) twice.

Mass spectrometry analysis was performed at the Proteomics Core Facility of the Shanghai Center for Plant Stress Biology. Matching raw MS data to peptide sequences was performed using Mascot software with the annotated proteins from the *N. benthamiana* draft genome sequence v. 0.4.4, which was obtained from the International Solanaceae Genomics Project (SOL) (https://solgenomics.net/), and AtRLP44-GFP sequence.

### Quantitative Real-Time PCR

For RNA analysis, a maximum of 100 mg of frozen *A. thaliana* seedling material was ground in a 2 ml reaction tube with the aid of a pre-cooled tissue lyser (TissuelyserII, Qiagen). RNA from ground tissue was extracted with an RNA purification Kit (Roboklon), following the manufacturer’s instructions. Synthesis of cDNA was carried out using AMV Reverse Trascriptase (EURx) following the manufacturer’s instructions. The cDNA reaction was diluted 1:10 in water and used for qPCR analysis with primers directed against BR marker genes *EXPA8* and *DWF4* or against *RLP44*.The SYBRR Green I nucleic acid gel stain (Sigma-Aldrich) was used for detection, *CLATHRIN*(At1g10730) was used as reference gene. qPCR reactions were run in a Rotor-Gene Q 2plex (Qiagen) and the amplification data extracted by the 75 Rotor-Gene Q 2plex software and analysed according to the method by (Muller et al., 2002). For primers, see Table S2. To assess *RLP44* transcript levels, PMEIox or *cnu2* seedlings transformed with untagged RLP44 constructs were ground in liquid nitrogen and RNA purification was performed with the OMEGA BIOTEK kit (OMEGA BIOTEK, Norcross, GA, USA) following the manufacturer’s instructions. DNAase treatment and cDNA synthesis were done with the iSript gDNA clear cDNA synthesis kit (BIO-RAD, Hercules, CA, USA). Quantitative reverse transcription-polymerase chain reaction (RT-PCR) was performed using *RLP44* primers listed in Supplementary Table S2 and *ACTIN (ACT2)* as reference gene.

### Genotyping

Presence of the *rlp44^cnu2^* mutation was assessed by CAPS marker using primers SW503 and SW504 and subsequent *HinfI* digestion. For the genotyping of the *bri1-null* T-DNA insertion, primers SW1378 and SW1379 were used for detection of the wild type allele. Presence of the T-DNA insertion was assessed with primers SW1377 and SW1379. For genotyping *pskr1-3*,primers SW1745 and SW1746 were used to assess presence of the wild type allele, and SW130 and SW1746 to assess presence of the T-DNA insertion. Primers SW1984 and SW1985 were used to detect the presence of the *PSKR2* wild type allele, and SW230 and SW1985 were used to detect the T-DNA. Presence of Gateway insertions was checked by using primers SW905 and SW906 directed against the attB1 and attB2 sequences, respectively. Presence of GreenGate insertions was assessed by PCR with primers SW1202 and SW1137.

### Xylem cell number analysis

Basic fuchsin staining of five-day-old seedling roots and CLSM analysis was performed as described (Holzwart et al., 2018).

### Microscopy

CLSM of Arabidopsis roots and *N. benthamiana* leaf discs was performed on a TCS SP5 II inverted Confocal Laser Scanning Microscope (Leica) or a LSM 510 Meta Confocal Laser Scanning Microscope (Zeiss). In the first case, a HCX PL APO lambda blue 63.0×1.20 water immersion objective (Leica) was used. In the second case, a Plan-Neofluoar 5.0×1.05, Plan-Neofluar 25.0×0.80 water immersion and C-Aprochromat 63.0×1.20 water immersion objectives were used. Excitation wavelength was set to 488 nm for GFP, 514 for YFP, and 561 nm for RFP or mCherry. Emission was recorded at 500-545 nm for GFP, at 545-573 nm for YFP and 620-670 nm for RFP or mCherry using HyD hybrid detectors (Leica) or photomultipliers (PMT) detectors (Zeiss). For inhibitor treatments, 6-day-old seedlings were incubated in 12-well plates using half-strength liquid MS, pH 5.8 supplemented with 20 μM Wortmannin (WM) (Sigma-Aldrich) or 50 μM of brefeldin A (BFA) (Sigma-Aldrich). For mock treatment, an equivalent volume of DMSO (Sigma-Aldrich) was used. Incubation with inhibitors took place at 22 °C in the dark for 165 min (WM) or 120 min (BFA) before imaging. FM4-64 staining was performed in half-strength liquid MS, pH 5.8 with 1 μM FM4-64 (Molecular probes) for 20 min. Seedlings were imaged with CLSM using 561 nm laser line for excitation and 670-750 nm range for emission detection. For quantification of oblique cell walls in root meristems, seedlings were grown on half-strength MS pates for five days, fixed in 4% paraformaldehyde solution, washed in PBS twice and incubated in ClearSee solution overnight (Kurihara et al., 2015; Ursache et al., 2018). Seedlings were stained for 60 min with 0.1% (w/v) Calcofluor White (Sigma-Aldrich) in ClearSee solution and subsequently washed for 30 min in ClearSee (Ursache et al., 2018). Seedlings were imaged using 405 nm laser line for excitation and 425-475 nm range for emission detection.

## Acknowledgements

We would like to thank Yi Wu and Pengcheng Wang for advice and technical help with proteomics, Clara Sanchez Rodriguez for sharing seeds of the amiTPL line; Heike Steininger for technical assistance; Falco Krüger and the Schumacher lab for help with microscopy; and Klaus Harter, Karin Schumacher, Thomas Greb, and Jan Lohmann for discussion. Mass spectrometry analysis was performed at the Proteomics Core Facility of the Shanghai Center for Plant Stress Biology. Research in the Wolf lab was supported by the German Research Foundation (DFG) through grants WO 1660/2-1 and SW 1660/6-2. SW was supported by the DFG through the Emmy Noether Programme.

## Author contributions

B.G.G., R.L-D., and S.W. designed the research. B.G.G. performed most of the experiments, E.H., C.S. and S.W. performed some experiments. All authors analysed data. B.G.G, R.L-D., and S.W. wrote the manuscript.

## References

Albert, M., Jehle, A.K., Fürst, U., Chinchilla, D., Boller, T., and Felix, G. (2013). A Two-Hybrid-Receptor Assay Demonstrates Heteromer Formation as Switch-On for Plant Immune Receptors. Plant Physiology 163: 1504–1509.

Bakker J, Spits M, Neefjes J, Berlin I (2017) The EGFR odyssey - from activation to destruction in space and time. J Cell Sci 130: 4087–4096

Belkhadir Y, Jaillais Y (2015) The molecular circuitry of brassinosteroid signaling. New Phytol 206: 522–540

Beltrao P, Albanese V, Kenner LR, Swaney DL, Burlingame A, Villen J, Lim WA, Fraser JS, Frydman J, Krogan NJ (2012) Systematic functional prioritization of protein posttranslational modifications. Cell 150: 413–425

Bowman, J.L. et al. (2017). Insights into Land Plant Evolution Garnered from the Marchantia polymorpha Genome. Cell 171: 287–304.e15.

Brutus A, Sicilia F, Macone A, Cervone F, De Lorenzo G (2010) A domain swap approach reveals a role of the plant wall-associated kinase 1 (WAK1) as a receptor of oligogalacturonides. Proc Natl Acad Sci U S A 107: 9452–9457

Bucherl CA, Jarsch IK, Schudoma C, Segonzac C, Mbengue M, Robatzek S, MacLean D, Ott T, Zipfel C (2017) Plant immune and growth receptors share common signalling components but localise to distinct plasma membrane nanodomains. Elife 6

Cheon J, Fujioka S, Dilkes BP, Choe S (2013) Brassinosteroids regulate plant growth through distinct signaling pathways in Selaginella and Arabidopsis. PLoS One 8: e81938

Couto D, Niebergall R, Liang X, Bucherl CA, Sklenar J, Macho AP, Ntoukakis V, Derbyshire P, Altenbach D, Maclean D, Robatzek S, Uhrig J, Menke F, Zhou JM, Zipfel C (2016) The Arabidopsis Protein Phosphatase PP2C38 Negatively Regulates the Central Immune Kinase BIK1. PLoS Pathog 12: e1005811

Couto D, Zipfel C (2016) Regulation of pattern recognition receptor signalling in plants. Nat Rev Immunol 16: 537–552

Dettmer J, Hong-Hermesdorf A, Stierhof YD, Schumacher K (2006) Vacuolar H+-ATPase activity is required for endocytic and secretory trafficking in Arabidopsis. Plant Cell 18: 715–730

Di Rubbo S, Irani NG, Kim SY, Xu ZY, Gadeyne A, Dejonghe W, Vanhoutte I, Persiau G, Eeckhout D, Simon S, Song K, Kleine-Vehn J, Friml J, De Jaeger G, Van Damme D, Hwang I, Russinova E (2013) The clathrin adaptor complex AP-2 mediates endocytosis of brassinosteroid insensitive1 in Arabidopsis. Plant Cell 25: 2986–2997

Domozych DS, Sorensen I, Popper ZA, Ochs J, Andreas A, Fangel JU, Pielach A, Sacks C, Brechka H, Ruisi-Besares P, Willats WG, Rose JK (2014) Pectin metabolism and assembly in the cell wall of the charophyte green alga Penium margaritaceum. Plant Physiol 165: 105–118

Dubeaux G, Vert G (2017) Zooming into plant ubiquitin-mediated endocytosis. Curr Opin Plant Biol 40: 56–62

Dubeaux G, Zelazny E, Vert G (2015) Getting to the root of plant iron uptake and cell-cell transport: Polarity matters! Commun Integr Biol 8: e1038441

Emans N, Zimmermann S, Fischer R (2002) Uptake of a fluorescent marker in plant cells is sensitive to brefeldin A and wortmannin. Plant Cell 14: 71–86

Erwig J, Ghareeb H, Kopischke M, Hacke R, Matei A, Petutschnig E, Lipka V (2017) Chitin-induced and CHITIN ELICITOR RECEPTOR KINASE1 (CERK1) phosphorylation-dependent endocytosis of Arabidopsis thaliana LYSIN MOTIFCONTAINING RECEPTOR-LIKE KINASE5 (LYK5). New Phytol 215: 382–396

Friedrichsen DM, Joazeiro CA, Li J, Hunter T, Chory J (2000) Brassinosteroid-insensitive-1 is a ubiquitously expressed leucine-rich repeat receptor serine/threonine kinase. Plant Physiol 123: 1247–1256

Fritz-Laylin LK, Krishnamurthy N, Tor M, Sjolander KV, Jones JD (2005) Phylogenomic analysis of the receptor-like proteins of rice and Arabidopsis. Plant Physiol 138: 611–623

Gadeyne A, Sanchez-Rodriguez C, Vanneste S, Di Rubbo S, Zauber H, Vanneste K, Van Leene J, De Winne N, Eeckhout D, Persiau G, Van De Slijke E, Cannoot B, Vercruysse L, Mayers JR, Adamowski M, Kania U, Ehrlich M, Schweighofer A, Ketelaar T, Maere S, Bednarek SY, Friml J, Gevaert K, Witters E, Russinova E, Persson S, De Jaeger G, Van Damme D (2014) The TPLATE adaptor complex drives clathrin-mediated endocytosis in plants. Cell 156: 691–704

Galindo-Trigo S, Blumke P, Simon R, Butenko MA (2020) Emerging mechanisms to fine-tune receptor kinase signaling specificity. Curr Opin Plant Biol 57: 41–51

Geldner N, Anders N, Wolters H, Keicher J, Kornberger W, Muller P, Delbarre A, Ueda T, Nakano A, Jurgens G (2003) The Arabidopsis GNOM ARF-GEF mediates endosomal recycling, auxin transport, and auxin-dependent plant growth. Cell 112: 219–230

Geldner N, Jurgens G (2006) Endocytosis in signalling and development. Curr Opin Plant Biol 9: 589–594

Grebe M, Xu J, Mobius W, Ueda T, Nakano A, Geuze HJ, Rook MB, Scheres B (2003) Arabidopsis sterol endocytosis involves actin-mediated trafficking via ARA6-positive early endosomes. Curr Biol 13: 1378–1387

Gronnier J, Legrand A, Loquet A, Habenstein B, Germain V, Mongrand S (2019) Mechanisms governing subcompartmentalization of biological membranes. Curr Opin Plant Biol 52: 114–123

Gust AA, Felix G (2014) Receptor like proteins associate with SOBIR1-type of adaptors to form bimolecular receptor kinases. Curr Opin Plant Biol 21C: 104–111

Haglund K, Dikic I (2012) The role of ubiquitylation in receptor endocytosis and endosomal sorting. J Cell Sci 125: 265–275

Hartmann J, Stuhrwohldt N, Dahlke RI, Sauter M (2013) Phytosulfokine control of growth occurs in the epidermis, is likely to be non-cell autonomous and is dependent on brassinosteroids. Plant J 73: 579–590

He Z, Wang ZY, Li J, Zhu Q, Lamb C, Ronald P, Chory J (2000) Perception of brassinosteroids by the extracellular domain of the receptor kinase BRI1. Science 288: 2360–2363

Hohmann U, Lau K, Hothorn M (2017) The Structural Basis of Ligand Perception and Signal Activation by Receptor Kinases. Annu Rev Plant Biol 68: 109–137

Hohmann U, Santiago J, Nicolet J, Olsson V, Spiga FM, Hothorn LA, Butenko MA, Hothorn M (2018) Mechanistic basis for the activation of plant membrane receptor kinases by SERK-family coreceptors. Proc Natl Acad Sci U S A 115: 3488–3493

Holt LJ, Tuch BB, Villen J, Johnson AD, Gygi SP, Morgan DO (2009) Global analysis of Cdk1 substrate phosphorylation sites provides insights into evolution. Science 325: 1682–1686

Holzwart E, Huerta AI, Glockner N, Garnelo Gomez B, Wanke F, Augustin S, Askani JC, Schurholz AK, Harter K, Wolf S (2018) BRI1 controls vascular cell fate in the Arabidopsis root through RLP44 and phytosulfokine signaling. Proc Natl Acad Sci U S A

Holzwart E, Wanke F, Glockner N, Hofte H, Harter K, Wolf S (2020) A mutant allele uncouples the brassinosteroid-dependent and independent functions of BRASSINOSTEROID INSENSITIVE 1. Plant Physiol 182: 669–678

Howe CL, Mobley WC (2004) Signaling endosome hypothesis: A cellular mechanism for long distance communication. J Neurobiol 58: 207–216

Jaillais Y, Belkhadir Y, Balsemao-Pires E, Dangl JL, Chory J (2011a) Extracellular leucine-rich repeats as a platform for receptor/coreceptor complex formation. Proc Natl Acad Sci U S A 108: 8503–8507

Jaillais Y, Hothorn M, Belkhadir Y, Dabi T, Nimchuk ZL, Meyerowitz EM, Chory J (2011b) Tyrosine phosphorylation controls brassinosteroid receptor activation by triggering membrane release of its kinase inhibitor. Genes Dev 25: 232–237

Jaillais Y, Ott T (2019) The nanoscale organization of the plasma membrane and its importance in signaling - a proteolipid perspective. Plant Physiol

Jamieson PA, Shan L, He P (2018) Plant cell surface molecular cypher: Receptor-like proteins and their roles in immunity and development. Plant Sci 274: 242–251

Karimi M, Inze D, Depicker A (2002) GATEWAY vectors for Agrobacterium-mediated plant transformation. Trends Plant Sci 7: 193–195

Kurihara D, Mizuta Y, Sato Y, Higashiyama T (2015) ClearSee: a rapid optical clearing reagent for whole-plant fluorescence imaging. Development 142: 4168–4179

Kutschmar A, Rzewuski G, Stuhrwohldt N, Beemster GT, Inze D, Sauter M (2009) PSK-alpha promotes root growth in Arabidopsis. New Phytol 181: 820–831

Ladwig F, Dahlke RI, Stuhrwohldt N, Hartmann J, Harter K, Sauter M (2015) Phytosulfokine Regulates Growth in Arabidopsis through a Response Module at the Plasma Membrane That Includes CYCLIC NUCLEOTIDE-GATED CHANNEL17, H+-ATPase, and BAK1. Plant Cell 27: 1718–1729

Lampropoulos A, Sutikovic Z, Wenzl C, Maegele I, Lohmann JU, Forner J (2013) GreenGate--a novel, versatile, and efficient cloning system for plant transgenesis. PLoS One 8: e83043

Li J, Chory J (1997) A putative leucine-rich repeat receptor kinase involved in brassinosteroid signal transduction. Cell 90: 929–938

Li J, Wen J, Lease KA, Doke JT, Tax FE, Walker JC (2002) BAK1, an Arabidopsis LRR receptor-like protein kinase, interacts with BRI1 and modulates brassinosteroid signaling. Cell 110: 213–222

Lin W, Lu D, Gao X, Jiang S, Ma X, Wang Z, Mengiste T, He P, Shan L (2013) Inverse modulation of plant immune and brassinosteroid signaling pathways by the receptorlike cytoplasmic kinase BIK1. Proc Natl Acad Sci U S A 110: 12114–12119

Liu D, Kumar R, Claus LAN, Johnson AJ, Siao W, Vanhoutte I, Wang P, Bender KW, Yperman K, Martins S, Zhao X, Vert G, Van Damme D, Friml J, Russinova E (2020) Endocytosis of BRASSINOSTEROID INSENSITIVE1 Is Partly Driven by a Canonical Tyr-Based Motif. Plant Cell 32: 3598–3612

Lu D, Lin W, Gao X, Wu S, Cheng C, Avila J, Heese A, Devarenne TP, He P, Shan L (2011) Direct ubiquitination of pattern recognition receptor FLS2 attenuates plant innate immunity. Science 332: 1439–1442

Luo Y, Scholl S, Doering A, Zhang Y, Irani NG, Rubbo SD, Neumetzler L, Krishnamoorthy P, Van Houtte I, Mylle E, Bischoff V, Vernhettes S, Winne J, Friml J, Stierhof YD, Schumacher K, Persson S, Russinova E (2015) V-ATPase activity in the TGN/EE is required for exocytosis and recycling in Arabidopsis. Nat Plants 1: 15094

Ma X, Xu G, He P, Shan L (2016) SERKing Coreceptors for Receptors. Trends Plant Sci

Martins S, Dohmann EM, Cayrel A, Johnson A, Fischer W, Pojer F, Satiat-Jeunemaitre B, Jaillais Y, Chory J, Geldner N, Vert G (2015) Internalization and vacuolar targeting of the brassinosteroid hormone receptor BRI1 are regulated by ubiquitination. Nat Commun 6: 6151

Matsubayashi Y, Ogawa M, Kihara H, Niwa M, Sakagami Y (2006) Disruption and overexpression of Arabidopsis phytosulfokine receptor gene affects cellular longevity and potential for growth. Plant Physiol 142: 45–53

Matsubayashi Y, Ogawa M, Morita A, Sakagami Y (2002) An LRR receptor kinase involved in perception of a peptide plant hormone, phytosulfokine. Science 296: 1470–1472

Mbengue M, Bourdais G, Gervasi F, Beck M, Zhou J, Spallek T, Bartels S, Boller T, Ueda T, Kuhn H, Robatzek S (2016) Clathrin-dependent endocytosis is required for immunity mediated by pattern recognition receptor kinases. Proc Natl Acad Sci U S A 113: 11034–11039

McKenna JF, Rolfe DJ, Webb SED, Tolmie AF, Botchway SW, Martin-Fernandez ML, Hawes C, Runions J (2019) The cell wall regulates dynamics and size of plasma-membrane nanodomains in Arabidopsis. Proc Natl Acad Sci U S A

Mithoe SC, Menke FL (2018) Regulation of pattern recognition receptor signalling by phosphorylation and ubiquitination. Curr Opin Plant Biol 45: 162–170

Monaghan J, Matschi S, Shorinola O, Rovenich H, Matei A, Segonzac C, Malinovsky FG, Rathjen JP, MacLean D, Romeis T, Zipfel C (2014) The Calcium-Dependent Protein Kinase CPK28 Buffers Plant Immunity and Regulates BIK1 Turnover. Cell Host Microbe 16: 605–615

Muller PY, Janovjak H, Miserez AR, Dobbie Z (2002) Processing of gene expression data generated by quantitative real-time RT-PCR. Biotechniques 32: 1372–1374, 1376, 1378–1379

Nam KH, Li J (2002) BRI1/BAK1, a receptor kinase pair mediating brassinosteroid signaling. Cell 110: 203–212

Nimchuk ZL, Tarr PT, Ohno C, Qu X, Meyerowitz EM (2011) Plant stem cell signaling involves ligand-dependent trafficking of the CLAVATA1 receptor kinase. Curr Biol 21: 345–352

Nishi H, Hashimoto K, Panchenko AR (2011) Phosphorylation in protein-protein binding: effect on stability and function. Structure 19: 1807–1815

Nishiyama T et al., (2018) The Chara Genome: Secondary Complexity and Implications for Plant Terrestrialization. Cell 174: 448–464 e424

Offringa R, Huang F (2013) Phosphorylation-dependent trafficking of plasma membrane proteins in animal and plant cells. J Integr Plant Biol 55: 789–808

Ortiz-Morea FA, Savatin DV, Dejonghe W, Kumar R, Luo Y, Adamowski M, Van den Begin J, Dressano K, Pereira de Oliveira G, Zhao X, Lu Q, Madder A, Friml J, Scherer de Moura D, Russinova E (2016) Danger-associated peptide signaling in Arabidopsis requires clathrin. Proc Natl Acad Sci U S A 113: 11028–11033

Park CJ, Peng Y, Chen X, Dardick C, Ruan D, Bart R, Canlas PE, Ronald PC (2008) Rice XB15, a protein phosphatase 2C, negatively regulates cell death and XA21-mediated innate immunity. PLoS Biol 6: e231

Pearlman SM, Serber Z, Ferrell JE, Jr. (2011) A mechanism for the evolution of phosphorylation sites. Cell 147: 934–946

Perraki A, DeFalco TA, Derbyshire P, Avila J, Sere D, Sklenar J, Qi X, Stransfeld L, Schwessinger B, Kadota Y, Macho AP, Jiang S, Couto D, Torii KU, Menke FLH, Zipfel C (2018) Phosphocode-dependent functional dichotomy of a common coreceptor in plant signalling. Nature 561: 248–252

Popper ZA, Michel G, Herve C, Domozych DS, Willats WG, Tuohy MG, Kloareg B, Stengel DB (2011) Evolution and diversity of plant cell walls: from algae to flowering plants. Annu Rev Plant Biol 62: 567–590

Proseus TE, Boyer JS (2006) Calcium pectate chemistry controls growth rate of Chara corallina. J Exp Bot 57: 3989–4002

Riano-Pachon DM, Kleessen S, Neigenfind J, Durek P, Weber E, Engelsberger WR, Walther D, Selbig J, Schulze WX, Kersten B (2010) Proteome-wide survey of phosphorylation patterns affected by nuclear DNA polymorphisms in Arabidopsis thaliana. BMC Genomics 11: 411

Richter S, Kientz M, Brumm S, Nielsen ME, Park M, Gavidia R, Krause C, Voss U, Beckmann H, Mayer U, Stierhof YD, Jurgens G (2014) Delivery of endocytosed proteins to the cell-division plane requires change of pathway from recycling to secretion. Elife 3: e02131

Robatzek S, Chinchilla D, Boller T (2006) Ligand-induced endocytosis of the pattern recognition receptor FLS2 in Arabidopsis. Genes Dev 20: 537–542

Sauter M (2015) Phytosulfokine peptide signalling. J Exp Bot 66: 5161–5169

Shiu SH, Bleecker AB (2001) Receptor-like kinases from Arabidopsis form a monophyletic gene family related to animal receptor kinases. Proc Natl Acad Sci U S A 98: 10763–10768

Sigismund S, Algisi V, Nappo G, Conte A, Pascolutti R, Cuomo A, Bonaldi T, Argenzio E, Verhoef LG, Maspero E, Bianchi F, Capuani F, Ciliberto A, Polo S, Di Fiore PP (2013) Threshold-controlled ubiquitination of the EGFR directs receptor fate. EMBO J 32: 2140–2157

Sigismund S, Confalonieri S, Ciliberto A, Polo S, Scita G, Di Fiore PP (2012) Endocytosis and signaling: cell logistics shape the eukaryotic cell plan. Physiol Rev 92: 273–366

Singh AP, Savaldi-Goldstein S (2015) Growth control: brassinosteroid activity gets context. J Exp Bot 66: 1123–1132

Smakowska-Luzan E, Mott GA, Parys K, Stegmann M, Howton TC, Layeghifard M, Neuhold J, Lehner A, Kong J, Grunwald K, Weinberger N, Satbhai Sb, Mayer D, Busch W, Madalinski M, Stolt-Bergner P, Provart NJ, Mukhtar MS, Zipfel C, Desveaux D, Guttman DS, Belkhadir Y (2018) An extracellular network of Arabidopsis leucine-rich repeat receptor kinases. Nature 553: 342–346

Song JH, Kwak SH, Nam KH, Schiefelbein J, Lee MM (2019) QUIRKY regulates root epidermal cell patterning through stabilizing SCRAMBLED to control CAPRICE movement in Arabidopsis. Nat Commun 10: 1744

Sun Y, Fan XY, Cao DM, Tang W, He K, Zhu JY, He JX, Bai MY, Zhu S, Oh E, Patil S, Kim TW, Ji H, Wong WH, Rhee SY, Wang ZY (2010) Integration of brassinosteroid signal transduction with the transcription network for plant growth regulation in Arabidopsis. Dev Cell 19: 765–777

Swaney DL, Beltrao P, Starita L, Guo A, Rush J, Fields S, Krogan NJ, Villen J (2013) Global analysis of phosphorylation and ubiquitylation cross-talk in protein degradation. Nat Methods 10: 676–682

Tyagi M, Shoemaker BA, Bryant SH, Panchenko AR (2009) Exploring functional roles of multibinding protein interfaces. Protein Sci 18: 1674–1683

Ursache, R., Andersen, T.G., Marhavý, P., and Geldner, N. (2018). A protocol for combining fluorescent proteins with histological stains for diverse cell wall components. Plant J 93: 399–412.

Van Damme D, Gadeyne A, Vanstraelen M, Inze D, Van Montagu MC, De Jaeger G, Russinova E, Geelen D (2011) Adaptin-like protein TPLATE and clathrin recruitment during plant somatic cytokinesis occurs via two distinct pathways. Proc Natl Acad Sci U S A 108: 615–620

Venne AS, Kollipara L, Zahedi RP (2014) The next level of complexity: crosstalk of posttranslational modifications. Proteomics 14: 513–524

Viotti C, Bubeck J, Stierhof YD, Krebs M, Langhans M, van den Berg W, van Dongen W, Richter S, Geldner N, Takano J, Jurgens G, de Vries SC, Robinson DG, Schumacher K (2010) Endocytic and secretory traffic in Arabidopsis merge in the trans-Golgi network/early endosome, an independent and highly dynamic organelle. Plant Cell 22: 1344–1357

Wang J, Cai Y, Miao Y, Lam SK, Jiang L (2009) Wortmannin induces homotypic fusion of plant prevacuolar compartments. J Exp Bot 60: 3075–3083

Wang J, Li H, Han Z, Zhang H, Wang T, Lin G, Chang J, Yang W, Chai J (2015) Allosteric receptor activation by the plant peptide hormone phytosulfokine. Nature 525: 265–268

Wolf S, Hematy K, Hofte H (2012a) Growth control and cell wall signaling in plants. Annu Rev Plant Biol 63: 381–407

Wolf S, Mravec J, Greiner S, Mouille G, Hofte H (2012b) Plant cell wall homeostasis is mediated by brassinosteroid feedback signaling. Curr Biol 22: 1732–1737

Wolf S, van der Does D, Ladwig F, Sticht C, Kolbeck A, Schurholz AK, Augustin S, Keinath N, Rausch T, Greiner S, Schumacher K, Harter K, Zipfel C, Hofte H (2014) A receptor-like protein mediates the response to pectin modification by activating brassinosteroid signaling. Proc Natl Acad Sci U S A 111: 15261–15266

Zhou J, Liu D, Wang P, Ma X, Lin W, Chen S, Mishev K, Lu D, Kumar R, Vanhoutte I, Meng X, He P, Russinova E, Shan L (2018) Regulation of Arabidopsis brassinosteroid receptor BRI1 endocytosis and degradation by plant U-box PUB12/PUB13-mediated ubiquitination. Proc Natl Acad Sci U S A 115: E1906–E1915

