## Supplemental Material for "Phosphorylation-dependent routing of RLP44 towards brassinosteroid or phytosulfokine signalling"

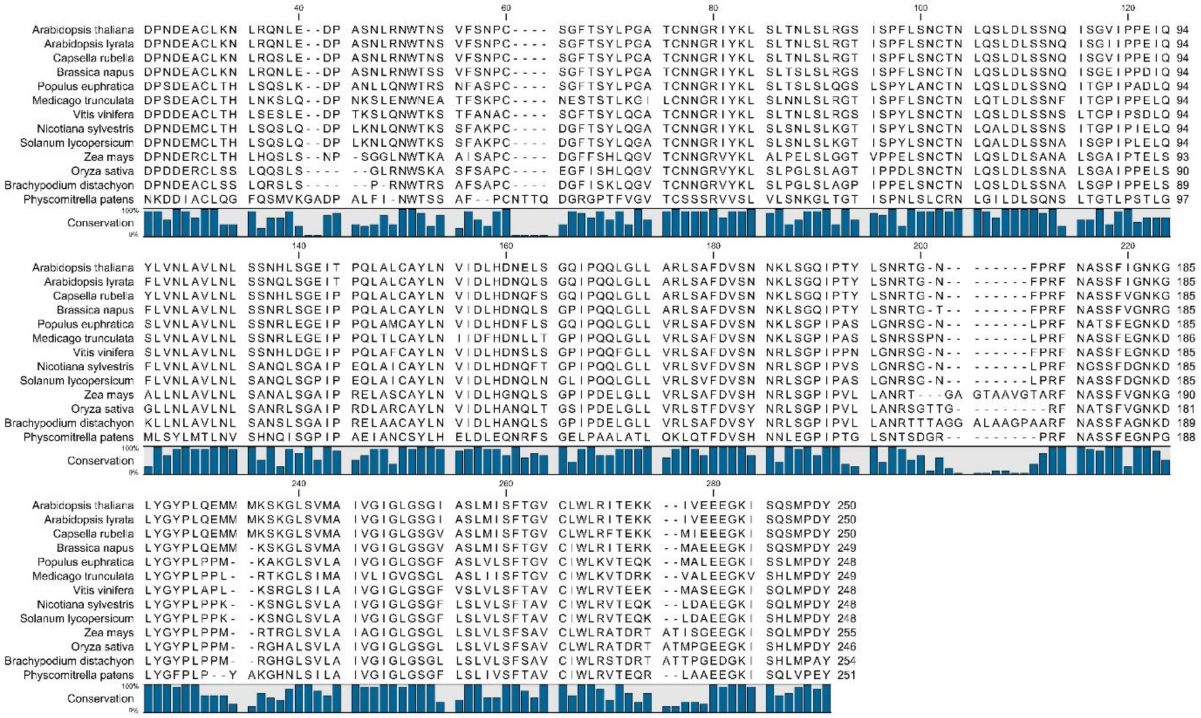

**Supplemental Figure S1.** Alignment of RLP44 amino acid sequences from various plant species.

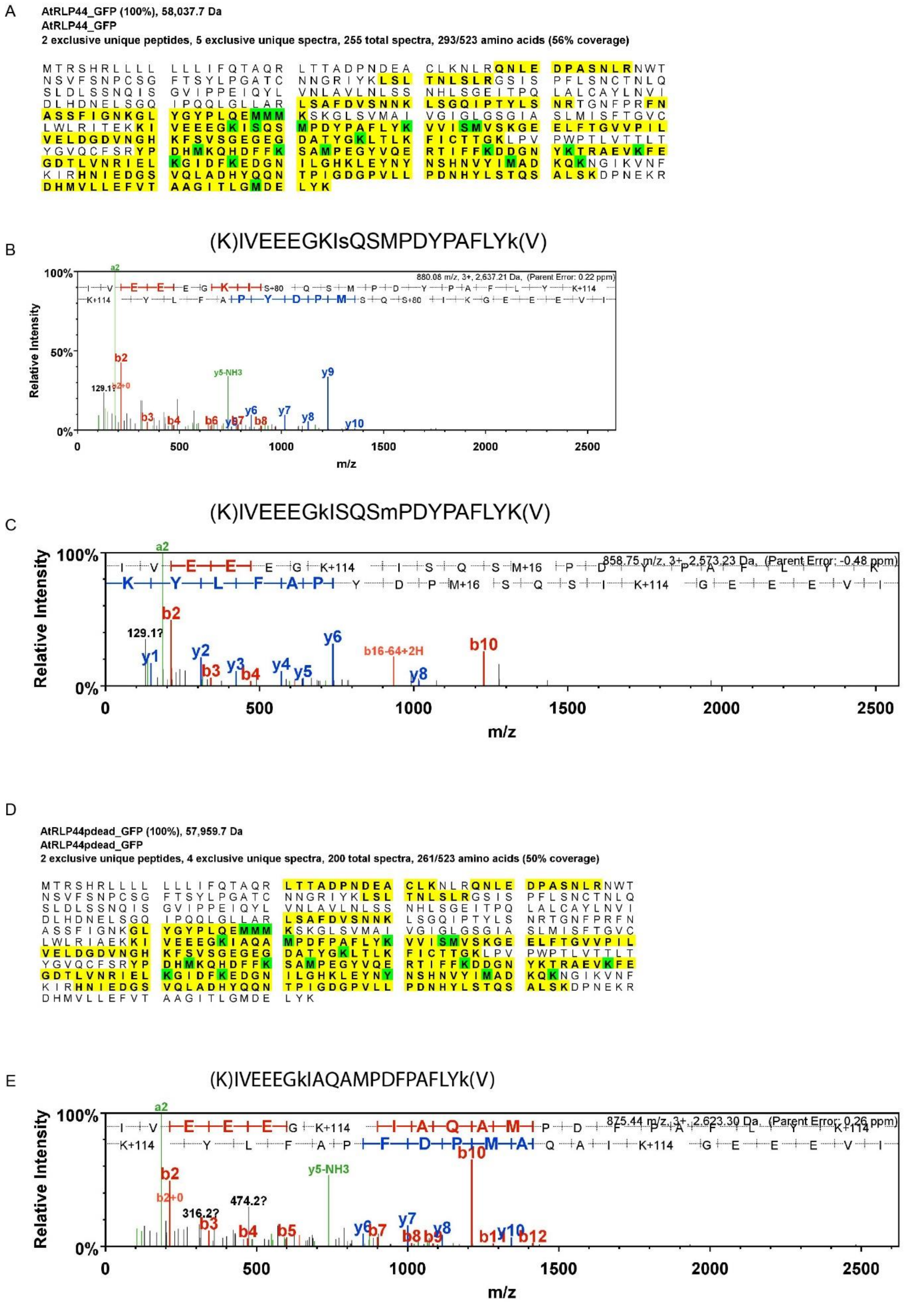

**Supplemental Figure S2.** Proteomics analysis of RLP44-GFP during transient expression in *N. benthamiana* and trypsin digestion. Peptide coverage (yellow) and modified amino acids (green) are indicated. Lower panes show spectra of peptides with phosphorylation of S268 and ubiquitination as indicated by double glycine remnant of K266. **(A)** RLP44-GFP WT peptide coverage. **(B)** RLP44-GFP WT peptide with phosphorylation of S268. **(C)** RLP44-GFP WT peptide with ubiquitination of K266. **(D)** RLP44-GFP Pdead peptide coverage. **(E)** RLP44-GFP Pdead peptide with ubiquitination of K266.

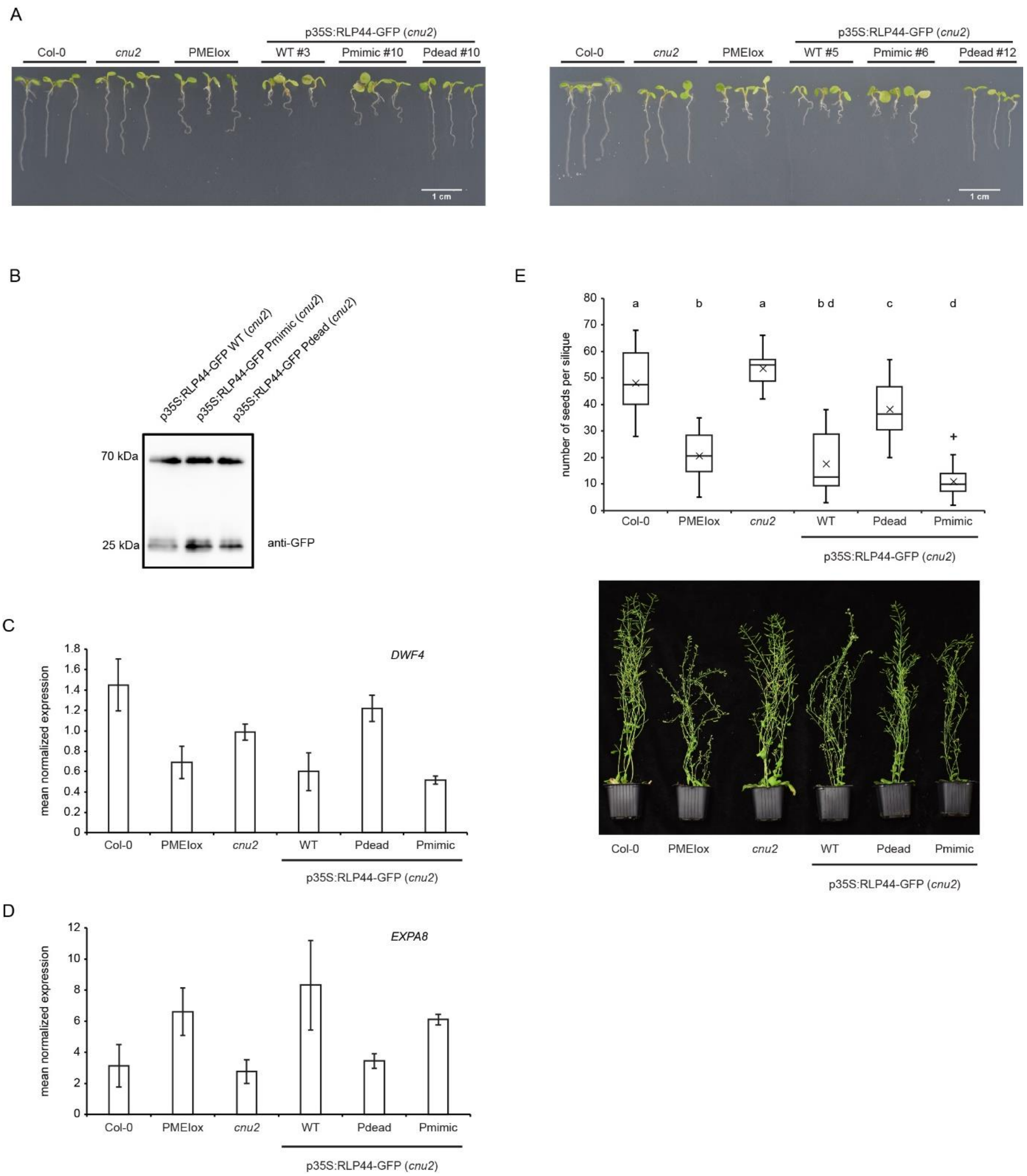

**Supplemental Figure S3.** Complementation of *cnu2* by RLP44-GFP WT and RLP44-GFP Pmimic, but not by RLP44-GFP Pdead. **(A)** Expression of RLP44-GFP WT and Pmimic, but not of Pdead, is able to complement the PMEIox suppressor mutant *cnu2* and leads to recovery of the PMEIox root waving phenotype in seedlings and contorted leaf arrangement in adult plants (see also Fig. 1). **(B)** Lines with comparable RLP44-GFP expression levels were selected based on Western Blot analysis with anti-GFP antiserum. **(C)** Quantitative Real-Time PCR analysis of the BR signalling marker gene *DWF4*. **(D)** Quantitative Real-Time PCR analysis of the BR signalling marker gene *EXPANSIN8*. **(E)** Analysis of seed yield in Col-0, PMEIox, *cnu2*, and *cnu2* complementation lines.

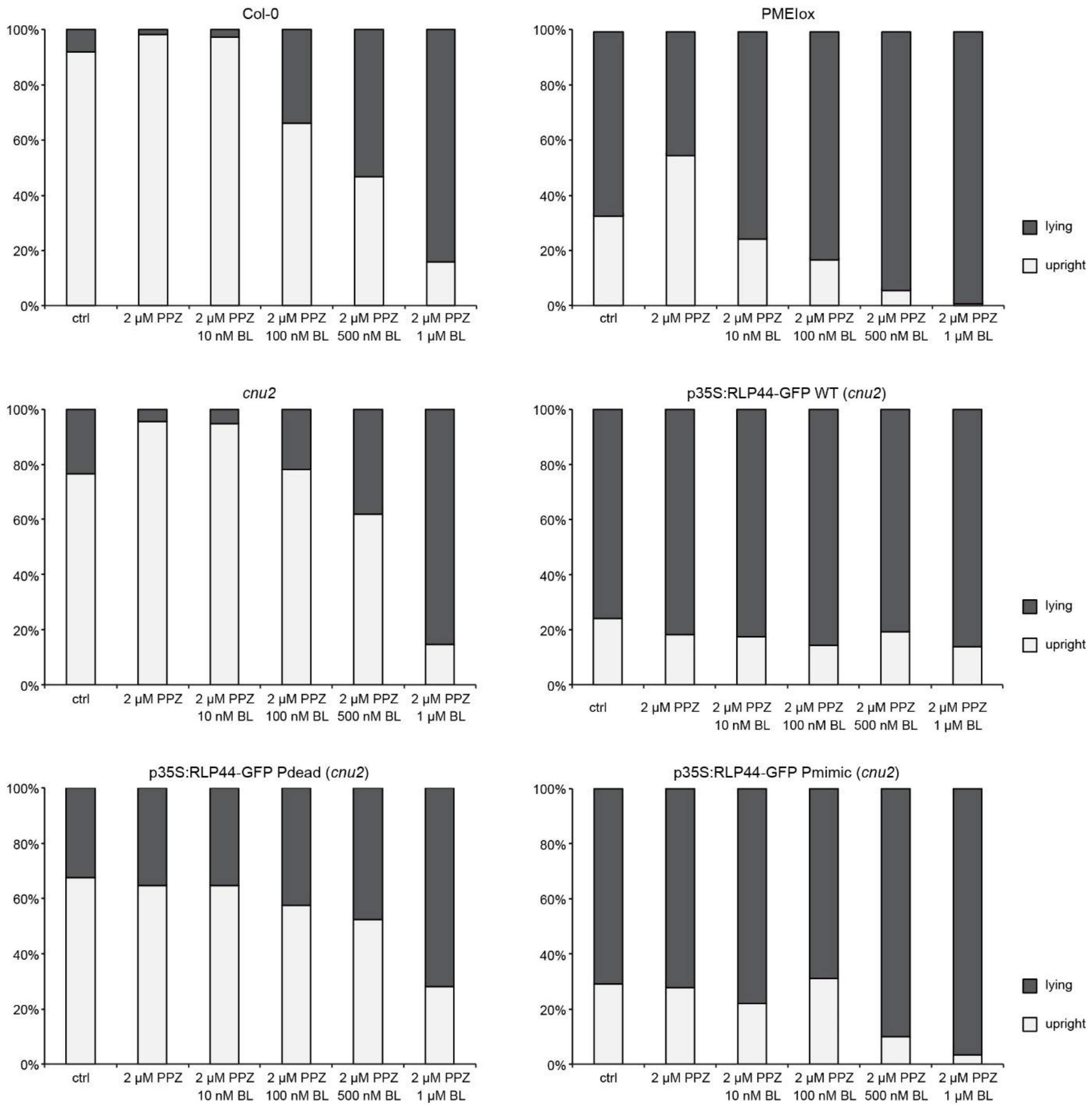

**Supplemental Figure S3.** RLP44-GFP WT and Pmimic, but not RLP44-GFP Pdead restore agravitropic growth of *cnu2* along the surface of agar plates in the dark as observed in PMEIox. Endogenous brassinosteroids were depleted with PPZ treatment to test sensitivity to increasing amounts of epi-brassinolide (BL)

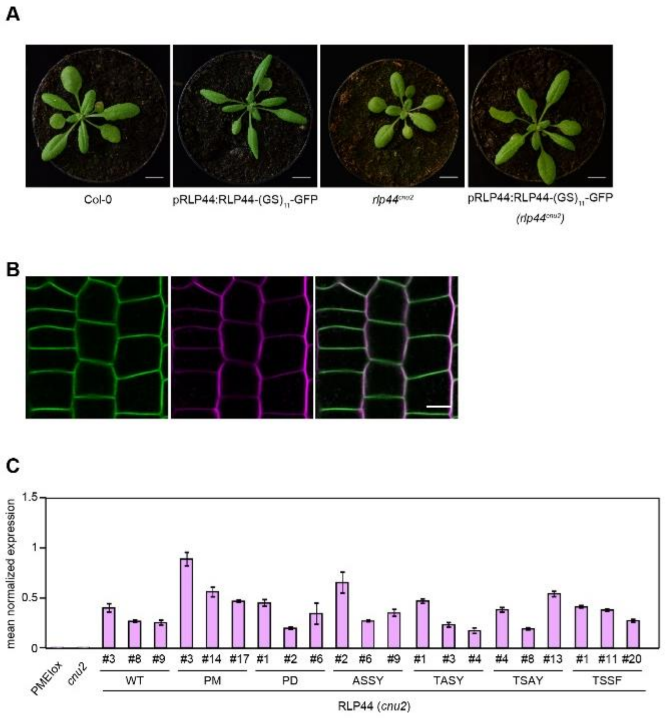

**Supplemental Figure S5.** Overall charge, rather than specific phosphosites modulate RLP44 function. **(A)** RLP44 overexpression phenotype (see Wolf et al., 2014) of hyperphosphorylated RLP44 under control of its own promoter. **(B)** Plasma membrane localization of hyperphosphorylated pRLP44:RLP44-(GS)$_{11}$-GFP. **(C)** Expression of *RLP44* in transgenic lines evaluated in Figure 10.

**Supplemental Table S1.** Mutants and transgenic lines used in this study.

| Mutant/transgenic line | |
|---|---|
| $rlp44^{cnu2}$ | Wolf *et al.*, 2014 |
| *cnu2* | Wolf *et al.*, 2014 |
| *bri1-null* | Jaillais *et al.*, 2011a |
| *pskr1-3 pskr2-1* | Kutschmar *et al.*, 2009 |
| *p35S:RLP44-GFP WT (cnu2)* | This study |
| *p35S:RLP44-GFP Pdead (cnu2)* | This study |
| *p35S:RLP44-GFP Pmimic (cnu2)* | This study |
| *p35S:RLP44-GFP WT* | This study |
| *p35S:RLP44-GFP Pdead* | This study |
| *p35S:RLP44-GFP Pmimic* | This study |
| *p35S:RLP44-GFP WT* ($rlp44^{cnu2}$) | This study |
| *p35S:RLP44-GFP Pdead* ($rlp44^{cnu2}$) | This study |
| *p35S:RLP44-GFP Pmimic* ($rlp44^{cnu2}$) | This study |
| *p35S:RLP44-GFP WT (bri1-null)* | Holzwart *et al.*, 2018 |
| *p35S:RLP44-GFP Pdead (bri1-null)* | This study |
| *p35S:RLP44-GFP Pmimic (bri1-null)* | This study |
| *p35S:RLP44-GFP WT (pskr1-3 pskr2-1)* | This study |
| *p35S:RLP44-GFP Pdead (pskr1-3 pskr2-1)* | This study |
| *pESTR:amiT-TML* | Gadeyne *et al.*, 2014 |
| *p35S:RLP44-GFP WT (pESTR:amiT-TML)* | This study |
| *p35S:RLP44-GFP Pdead (pESTR:amiT-TML)* | This study |
| *pRLP44:RLP44* WT (*cnu2*) | This study |
| *pRLP44:RLP44* Pdead (*cnu2)* | This study |
| *pRLP44:RLP44* Pmimic (*cnu2*) | This study |
| *pRLP44:RLP44 T256A (cnu2*) | This study |
| *pRLP44:RLP44 S268A (cnu2*) | This study |
| *pRLP44:RLP44 S270A (cnu2*) | This study |
| *pRLP44:RLP44 Y274F (cnu2*) | This study |
| *pRLP44:RLP44-(GS)11-GFP* | Holzwart et al., 2018 |
| *pRLP44:RLP44-(GS)11-GFP WT (cnu2)* | This study |
| *pRLP44:RLP44-(GS)11-GFP Pmimic (cnu2)* | This study |
| *pRLP44:RLP44-(GS)11-GFP (Pdead (cnu2)* | This study |

| *Primer No.* | *Primer name* | *Sequence (5′ → 3′)* | *target* |
|---|---|---|---|
| SW660 | RLP44_GW_L | GGGGACAAGTTTGTACAAAAAAGCAGGCTATGACAAGGAGTCACCGGTTAC | At3g49750 |
| SW670 | RLP_GW_WToY_R | GGGGACCACTTTGTACAAGAAAGCTGGGTtGTAATCAGGCATAGATTGAC | At3g49750 |
| SW666 | RLP44_SDMT256A_F | gtttatggttgaggattgctgagaagaagattgttg | At3g49750 |
| SW667 | RLP44_SDMT256A_R | caacaatcttcttctcagcaatcctcaaccataaac | At3g49750 |
| SW668 | RLP_S268,270A_Y274F_Rneu | GGGGACCACTTTGTACAAGAAAGCTGGGTagaaatcaggcatagcttgagcaatcttaccttcttcttcaac | At3g49750 |
| SW672 | RLP44_SDMT256E_F | GAGGATTGAAGAGAAGAAGATTGTTGAAGAAG | At3g49750 |
| SW673 | RLP44_SDMT256E_R | ATCTTCTTCTCTTCAATCCTCAACCATAAAC | At3g49750 |
| SW671 | RLP_SSY-EEE_R | GGGGACCACTTTGTACAAGAAAGCTGGGTagaaatcaggcatagattgactaatc | At3g49750 |
| SW661 | RLP44_Y274F_R | GGGGACCACTTTGTACAAGAAAGCTGGGTtttcatcaggcatttcttgttcaatcttaccttcttcttcaac | At3g49750 |
| SW662 | RLP44_S268A_R | GGGGACCACTTTGTACAAGAAAGCTGGGTagtaatcaggcatagcttgactaatcttaccttcttc | At3g49750 |
| SW663 | RLP44_S270A_R | GGGGACCACTTTGTACAAGAAAGCTGGGTagtaatcaggcatagattgagcaatcttaccttcttcttcaac | At3g49750 |
| SW2446 | RLP44-if-f | tgAagcttGGTCTCaGGCTcAATGACAAGGAGTCACCGG | At3g49750 |
| SW2447 | RLP44WT-if-r | atGGCACCCGCCCCTGCTCcGTAATCAGGCATAGATTG | At3g49750 |
| SW2448 | GAGAGA-GFP-if-f | gGAGCAGGGGCGGGTGCC | GFP |
| SW2449 | GAGAGA-GFP-if-r | cgaGAATTcGGTCTCaCTGAttacttgtacagctcgtcc | GFP |
| SW2450 | RLP44TS-AY -if-r | atGGCACCCGCCCCTGCTCcGTAATCAGGCATAGcTTG | At3g49750 |
| SW2452 | RLP44TS-SF-if-r | atGGCACCCGCCCCTGCTCcGaAATCAGGCATAGATTG | At3g49750 |
| SW2454 | RLP44AA-AF-if-r | atGGCACCCGCCCCTGCTCcGaAATCAGGCATAGcTTG | At3g49750 |
| SW2455 | RLP44EE-EE-if-r | atGGCACCCGCCCCTGCTCcttcATCAGGCATttcTTG | At3g49750 |
| SW3000 | RLP44prom_F | GGGGACAAGTTTGTACAAAAAAGCAGGCTTTTGCGATATTTTTGGCTGTC | At3g49750 |
| SW3001 | RLP44stop_R | GGGGACCACTTTGTACAAGAAAGCTGGGTTTTAGTAATCAGGCATAGATTGACT | At3g49750 |
| SW3002 | RLP44_TASY_stop_R | GGGGACCACTTTGTACAAGAAAGCTGGGTTTTAGTAATCAGGCATAGATTG | At3g49750 |
| SW3003 | RLP44_PM_stop_R | GGGGACCACTTTGTACAAGAAAGCTGGGTTTTATTCATCAGGCATTTCTTGTTCAATCTT | At3g49750 |
| SW3004 | RLP44_PD_stop_R | GGGGACCACTTTGTACAAGAAAGCTGGGTTTTAGAAATCAGGCATAGCTTGAGCAATCTT | At3g49750 |
| SW3005 | RLP44_TSAY_stop_R | GGGGACCACTTTGTACAAGAAAGCTGGGTTTTAGTAATCAGGCATAGCTTGACTAATCTT | At3g49750 |
| SW3006 | RLP44_TSSF_stop_R | GGGGACCACTTTGTACAAGAAAGCTGGGTTTTAGAAATCAGGCATAGATTGACTAATC | At3g49750 |
| SW1179 | RLP44_GG_F | AACAGGTCTCAGGCTCAATGACAAGGAGTCACCGGTTA | At3g49750 |
| SW1205 | RLP44_GG_R | AACAGGTCTCACTGAGTAATCAGGCATAGATTGAC | At3g49750 |
| SW1367 | RLP44_PD_GGC_R | AACAGGTCTCACTGAGAAATCAGGCATAGCTTG | At3g49750 |

| | | | |
|---|---|---|---|
| SW1368 | RLP44_PM_GGC_R | AACAGGTCTCACTGATTCATCAGGCATTTCTTG | At3g49750 |
| SW503 | RLP44-4_CAPS_F | AATCTACAAACTCTCACTCAC | At3g49750 |
| SW504 | RLP44-4_CAPS_R | CTGACCGGATAATTCGTTATC | At3g49750 |
| SW1377 | GK-O8409 | ATATTGACCATCATACTCATTGC | |
| SW1378 | GK-134E10_F | TAGCGGAAACAAAATCAGTGG | At4g39400 |
| SW1379 | GK-134E10_R | TCGTTCCATTGAAGAGATTGG | At4g39400 |
| SW1754 | pskr1-3_F | CTCGCTTTCTGGTATGACGAG | At2g02220 |
| SW1746 | pskr1-3_R | TCCGAAACTATACACATCGCC | At2g02220 |
| SW1984 | pskr2-1_F | TTCTTAGACTGTTTGGCTCGG | At5g53890 |
| SW1985 | pskr2-1_R | GCGTTACAAACATGCAACAAG | At5g53890 |
| SW230 | LBb1.3 | ATTTTGCCGATTTCGGAAC | |
| SW905 | attB1 | ACAAGTTTGTACAAAAAAGCAGGCT | |
| SW906 | attB2 | ACCACTTTGTACAAGAAAGCTGGGT | |
| SW1202 | pGG-Bdummy_F | GTATTCAGTCGACTGGTACCAAC | |
| SW1137 | pGGA/C000_R | CAGATTGTACTGAGAGTGCACC | |
| SW521 | AtEXP8_F | CCGAAATAACTAACCCCTCCTC | At2g40610 |
| SW522 | AtEXP8_R | TAGCCACAAGCTCCGCCCAT | At2g40610 |
| SW803 | DWF4_F | CAACAGCAAAACAACGGAGCG | At3g50660 |
| SW804 | DWF4_R | TCTGAACCAGCACATAGCCTTG | At3g50660 |
| SW1015 | Clath_F | TCGATTGCTTGGTTTGGAAGAT | At1g10730 |
| SW1016 | Clath_R | GCACTTAGCGTGGACTCTGTTTGC | At1g10730 |
| SW612 | RLP44COD2_F | TCAGATTCCGCAGCAATTAG | At3g49750 |
| SW613 | RLP44COD2_R | TCCTGCAACGGATAACCATA | At3g49750 |
| | ACT2_F | CAGTGTCTGGATCGGTGGTT | At3g18780 |
| | ACT2_R | TGAACGATTCCTGGACCTGC | At3g18780 |

**Supplemental Table S2.** Oligonucleotides used in this study.

| | | | | |
|---|---|---|---|---|
| **pSW362** | **pRLP44:RLP44-(GS)$_{11}$-GFP WT** | | | |
| | *Name* | *Internal name* | *Source* | *Primers* |
| “Promoter” module | pRLP44 | pSW299 | Holzwart et al., 2018 | |
| “N-tag” module | B-dummy | pGGB003 | Lampropoulos et al., 2013 | |
| “CDS” module | RLP44 | pSW334 | Holzwart et al., 2018 | SW1179-1205 |
| “C-tag” module | (GS)$_{11}$-GFP | PGGD001 | Lampropoulos et al., 2013 | |
| “Terminator” module | tUBQ10 | pGGE009 | Lampropoulos et al., 2013 | |
| “Resistance” module | SulfR | pGGF006 | Lampropoulos et al.,2013 | |
| Destination vector | | pGGZ0001 | Lampropoulos et al.,2013 | |
| **pSW567** | **pRLP44:RLP44-(GS)$_{11}$-GFP Pmimic** | | | |
| | *Name* | *Internal name* | *Source* | *Primers* |
| “Promoter” module | pRLP44 | pSW299 | Holzwart et al., 2018 | |
| “N-tag” module | B-dummy | pGGB003 | Lampropoulos et al., 2013 | |
| “CDS” module | RLP44pmimic | pSW519 | This study | SW1179-1368 |
| “C-tag” module | (GS)$_{11}$-GFP | PGGD001 | Lampropoulos et al., 2013 | |
| “Terminator” module | tUBQ10 | pGGE009 | Lampropoulos et al., 2013 | |
| “Resistance” module | SulfR | pGGF006 | Lampropoulos et al.,2013 | |
| Destination vector | | pGGZ0001 | Lampropoulos et al.,2013 | |
| **pSW566** | **pRLP44:RLP44-(GS)$_{11}$-GFP Pdead** | | | |
| | *Name* | *Internal name* | *Source* | *Primers* |
| “Promoter” module | pRLP44 | pSW299 | Holzwart et al., 2018 | |
| “N-tag” module | B-dummy | pGGB003 | Lampropoulos et al., 2013 | |
| “CDS” module | RLP44pdead | pSW518 | This study | SW1179-1367 |
| “C-tag” module | (GS)$_{11}$-GFP | PGGD001 | Lampropoulos et al., 2013 | |
| “Terminator” module | tUBQ10 | pGGE009 | Lampropoulos et al., 2013 | |
| “Resistance” module | SulfR | pGGF006 | Lampropoulos et al.,2013 | |
| Destination vector | | pGGZ0001 | Lampropoulos et al.,2013 | |
| **pSW1027** | **pRLP44:RLP44-GAGAGA-GFP WT** | | | |
| “Promoter” module | pRLP44 | pSW299 | Holzwart et al., 2018 | |
| “N-tag” module | B-dummy | pGGB003 | Lampropoulos et al., 2013 | |
| “CDS” module | RLP44-GAGAGA-GFP | pSW999 | This study | SW2446-2449 |
| “C-tag” module | D-dummy | pGGD002 | Lampropoulos et al., 2013 | |
| “Terminator” module | tUBQ10 | pGGE009 | Lampropoulos et al., 2013 | |
| “Resistance” module | SulfR | pGGF006 | Lampropoulos et al.,2013 | |
| Destination vector | | pGGZ0001 | Lampropoulos et al.,2013 | |
| **pSW1019** | **pRLP44:RLP44-GAGAGA-GFP Pmimic** | | | |
| “Promoter” module | pRLP44 | pSW299 | Holzwart et al., 2018 | |
| “N-tag” module | B-dummy | pGGB003 | Lampropoulos et al., 2013 | |
| “CDS” module | RLP44-GAGAGA-GFP Pmimic | pSW1018 | This study | SW2446-2455, 2448+2449 |
| “C-tag” module | D-dummy | pGGD002 | Lampropoulos et al., 2013 | |
| “Terminator” module | tUBQ10 | pGGE009 | Lampropoulos et al., 2013 | |
| “Resistance” module | SulfR | pGGF006 | Lampropoulos et al.,2013 | |
| Destination vector | | pGGZ0001 | Lampropoulos et al.,2013 | |
| **pSW1017** | **pRLP44:RLP44-GAGAGA-GFP Pdead** | | | |
| “Promoter” module | pRLP44 | pSW299 | Holzwart et al., 2018 | |
| “N-tag” module | B-dummy | pGGB003 | Lampropoulos et al., 2013 | |
| “CDS” module | RLP44-GAGAGA-GFP Pdead | pSW1007 | This study | SW2446+2454, 2448+2449 |

| | | | | |
|---|---|---|---|---|
| "C-tag" module | D-dummy | pGGD002 | Lampropoulos et al., 2013 | |
| "Terminator" module | tUBQ10 | pGGE009 | Lampropoulos et al., 2013 | |
| "Resistance" module | SulfR | pGGF006 | Lampropoulos et al.,2013 | |
| Destination vector | | pGGZ0001 | Lampropoulos et al.,2013 | |
| **pSW1026** | **pRLP44:RLP44-GAGAGA-GFP ASSY** | | | |
| "Promoter" module | pRLP44 | pSW299 | Holzwart et al., 2018 | |
| "N-tag" module | B-dummy | pGGB003 | Lampropoulos et al., 2013 | |
| "CDS" module | RLP44-GAGAGA-GFP ASSY | pSW1000 | This study | SW2446+2447, 2448+2449 |
| "C-tag" module | D-dummy | pGGD002 | Lampropoulos et al., 2013 | |
| "Terminator" module | tUBQ10 | pGGE009 | Lampropoulos et al., 2013 | |
| "Resistance" module | SulfR | pGGF006 | Lampropoulos et al.,2013 | |
| Destination vector | | pGGZ0001 | Lampropoulos et al.,2013 | |
| **pSW1028** | **pRLP44:RLP44-GAGAGA-GFP TASY** | | | |
| "Promoter" module | pRLP44 | pSW299 | Holzwart et al., 2018 | |
| "N-tag" module | B-dummy | pGGB003 | Lampropoulos et al., 2013 | |
| "CDS" module | RLP44-GAGAGA-GFP TASY | pSW1002 | This study | SW2446+2447, 2448+2449 |
| "C-tag" module | D-dummy | pGGD002 | Lampropoulos et al., 2013 | |
| "Terminator" module | tUBQ10 | pGGE009 | Lampropoulos et al., 2013 | |
| "Resistance" module | SulfR | pGGF006 | Lampropoulos et al.,2013 | |
| Destination vector | | pGGZ0001 | Lampropoulos et al.,2013 | |
| **pSW1014** | **pRLP44:RLP44-GAGAGA-GFP TSAY** | | | |
| "Promoter" module | pRLP44 | pSW299 | Holzwart et al., 2018 | |
| "N-tag" module | B-dummy | pGGB003 | Lampropoulos et al., 2013 | |
| "CDS" module | RLP44-GAGAGA-GFP TSAY | pSW1004 | This study | SW2446+2450, 2448+2449 |
| "C-tag" module | D-dummy | pGGD002 | Lampropoulos et al., 2013 | |
| "Terminator" module | tUBQ10 | pGGE009 | Lampropoulos et al., 2013 | |
| "Resistance" module | SulfR | pGGF006 | Lampropoulos et al.,2013 | |
| Destination vector | | pGGZ0001 | Lampropoulos et al.,2013 | |
| **pSW1015** | **pRLP44:RLP44-GAGAGA-GFP TSSF** | | | |
| "Promoter" module | pRLP44 | pSW299 | Holzwart et al., 2018 | |
| "N-tag" module | B-dummy | pGGB003 | Lampropoulos et al., 2013 | |
| "CDS" module | RLP44-GAGAGA-GFP TSSF | pSW1005 | This study | SW2446+2452, 2448+2449 |
| "C-tag" module | D-dummy | pGGD002 | Lampropoulos et al., 2013 | |
| "Terminator" module | tUBQ10 | pGGE009 | Lampropoulos et al., 2013 | |
| "Resistance" module | SulfR | pGGF006 | Lampropoulos et al.,2013 | |
| Destination vector | | pGGZ0001 | Lampropoulos et al.,2013 | |

**Supplemental Table S3.** Overview of constructs generated with GreenGate cloning.
